# Multimodal profiling of human sensory neurons links electrical properties to transcriptional identity

**DOI:** 10.1101/2025.03.25.645367

**Authors:** Jiwon Yi, Lite Yang, Allie J. Widman, Alexa Toliver, Zachariah Bertels, John Smith Del Rosario, Adam J. Dourson, Prashant Gupta, Rakesh Kumar, Jun-Nan Li, Maria Payne, Richard A. Slivicki, Juliet M. Mwirigi, Alexander Chamessian, Kesav Kothapalli, Grace Moore, John Lemen, Bryan A. Copits, Robert W. Gereau, PRECISION Human Pain Network

**Author notes:** Corresponding authors. Address: Department of Anesthesiology, Washington University Pain Center, Washington University School of Medicine, 660 South Euclid Avenue Campus Box 8054, St. Louis, 63110, MO, United States. (B.A. Copits) and (R.W. Gereau). URL: pain.wustl.edu. These authors contributed equally to this work (Zachariah Bertels, John Smith Del Rosario, Adam J. Dourson, Prashant Gupta, Rakesh Kumar, Jun-Nan Li, Maria Payne, Richard A. Slivicki).

## Abstract

Despite major advances in pain science, the approval of novel therapeutics has been slow. A major cause for the lack of new analgesics may be fundamental biological differences between humans and model organisms used in preclinical research. Large-scale transcriptional profiling efforts on human dorsal root ganglia (hDRG) have now identified at least 22 distinct neuronal subtypes; however, a significant knowledge gap exists in ascribing functional phenotypes to these diverse neuronal populations. In this study, we use Patch-seq recordings in hDRG to link electrical properties to transcriptionally defined cell types. First, through unbiased clustering of electrophysiological properties from 228 hDRG neurons, we identify three electrophysiological subtypes (E-types). Next, we show that E-types can be mapped onto specific transcriptional classes (T-types) of hDRG neurons. We find that donor’s pain history is associated with E-type-specific differences in electrical properties, some of which may be associated with higher expression of voltage-gated sodium channels, NaV1.7 and NaV1.8. These results highlight the importance of using multimodal profiling to better understand human sensory neuron biology and may help reveal novel therapeutic targets driving chronic pain.

**Teaser:** Mapping electrical properties to transcriptional profiles of human sensory neurons offers new insights into pain neurobiology

## Introduction

Chronic pain affects an estimated 20% of people worldwide, resulting in disability, reduced quality of life, and the dearth of treatment options is a major driver of the current opioid epidemic in the U.S.(*1–4*). While a multitude of potential new analgesic targets have been identified in preclinical studies in animal models, the vast majority of efforts to translate these findings into new treatments have failed in clinical trials. As such, available options for non-opioid pharmacotherapy in patients are still limited(*5*). While there are many possible reasons that translational efforts in pain fail so frequently, it is becoming increasingly apparent that a poor understanding of the relationship between human physiology and that of animals used in preclinical studies is a major contributor. Understanding the cellular identity and physiology of human sensory neurons and how these differ in the context of painful conditions is therefore critical to better translate preclinical findings from rodents to humans and to identify unique properties of human sensory neurons that can be targeted for therapeutic development. Collecting this type of information is the current goal of the NIH HEAL Initiative’s PRECISION human pain network (*6*). Here, we describe progress on one PRECISION network study where we seek to identify diverse electrophysiological phenotypes of human sensory neurons from the dorsal root ganglion (DRG), map these properties onto genetically identified cell types and understand differences in physiology and gene expression seen in the context of chronic pain.

Nociceptor hyperactivity following injury or inflammation is a critical driver in the development and/or maintenance of pain (*7*, *8*), and both C-fiber and A-fiber nociceptors display ectopic spontaneous activity and hyperexcitability in animal models of neuropathic, inflammatory, and chemotherapy-induced pain. The relevance of these findings is further bolstered by the observation that human dorsal root ganglia (hDRG) neurons from patients with neuropathic pain are much more likely to have spontaneous activity (*9–11*). Whether similar pain-associated hyperexcitability of hDRG neurons is seen in other pain conditions, such as musculoskeletal pain, is yet unknown. Interestingly, in a rat model of neuropathic pain, nociceptor sensitization is only seen in neurons that have non-accommodating (repetitively firing) spike patterns, but not in rapidly accommodating (single spiking) nociceptors (*12*). Similarly, bradykinin-induced hyperexcitability of hDRG neurons varies between repetitively firing and single spiking neurons (*13*). These findings suggest that certain electrophysiological features are associated with cell types that are sensitized in the context of pain-inducing injury or inflammation.

A wealth of studies in rodent DRG has revealed that nociceptors themselves are a heterogenous population that consist of multiple specialized sub-populations defined by distinct chemo- and mechano-sensitivities, transcriptional profiles, developmental regulation, and projection patterns (*14–25*). Molecularly defined subpopulations of mouse nociceptors display distinct intrinsic membrane properties, suggesting that transcriptional profiles and physiological properties are linked (*25*, *26*). While single-cell transcriptomics studies using hDRG show that core functional groups of sensory neurons appear conserved from rodents to humans, the relative proportion of different functional populations differ across species (*27–30*). Compared to rodent, hDRG neurons also more broadly express nociceptive receptors, ion channels, and neuropeptides (*31–37*), contain a higher proportion of chemosensitive neurons, and have different intrinsic membrane properties and spike kinetics (*38*). However, a multimodal characterization linking the transcriptional and physiological characteristics of human sensory neurons is currently lacking.

In this study, we address this gap by combining patch-clamp electrophysiology paired with single-cell RNA sequencing (Patch-seq (*39*)) in hDRG neurons obtained from postmortem organ donors. By profiling the excitability of over 200 hDRG neurons obtained from 56 donors, we identify three electrophysiological subtypes (E-types) of hDRG neurons that show distinct membrane properties and spike kinetics. We identify four transcriptional hDRG neuronal classes (T-types), which E-types can be broadly mapped onto. Next, we utilize sub-group analysis to show that hDRG neurons from donors with pain history demonstrate E-type dependent differences in spike kinetic properties and differential expression of ion channels. Together, these findings suggest that hDRG neurons show physiological specialization which may be associated with functional diversity and illustrate ways in which the large-scale collection of these types of multimodal data can generate insights into potential underlying causes of chronic pain to spur new therapeutic development.

## Results

### Unbiased clustering of small- to medium-size human somatosensory neurons based on physiological characteristics

Rodent sensory neurons can be organized into clusters that share similar physiological characteristics, and these clusters are associated with distinct transcriptional profiles and molecularly defined subpopulations of sensory neurons (*12*, *25*, *26*, *40*). Here, we tested whether such physiological heterogeneity also exists in hDRG neurons *in vitro*. Using primary culture of hDRG obtained from postmortem organ donors (**Table 1, Table S1**), we performed patch-clamp experiments to assess the membrane excitability of hDRG neurons. As prior work has demonstrated enrichment of human nociceptor populations among smaller diameter sensory neurons, whereas non-nociceptive subtypes like proprioceptors and low threshold mechanoreceptors are found in hDRG neurons with larger soma size (*33*, *34*) we focused our investigation on small- to medium-sized hDRG neurons (mean cell diameter ± SD: 40.65 ± 11.23µm). A small percentage of patch-clamp recordings (4.8%) were performed on larger hDRG neurons with diameter >60µm (**Fig. 1A**). To determine if different hDRG cell types exhibit distinct electrical properties, we extracted 10 electrophysiological features from 228 hDRG recordings (**Fig. 1B-D)**.

**Fig. 1.**
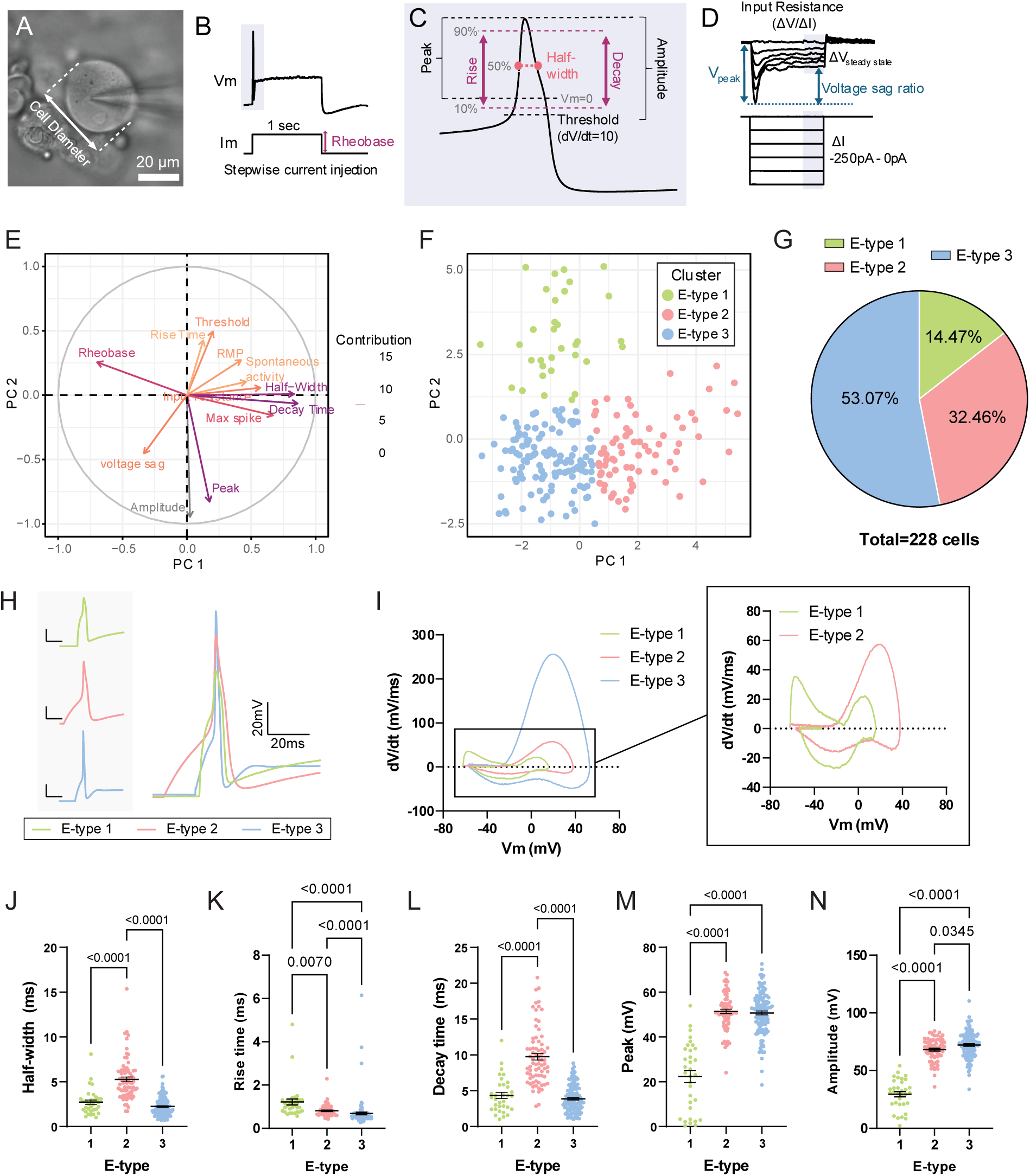
Human DRG exhibit three electrophysiologically defined cell types (E-types). (**A**) Image of cultured hDRG neuron recorded for this study. (**B**) Example action potential (AP) voltage trace (Vm, top) in response to a 1-second depolarizing current injection (Im, bottom) at threshold (rheobase). (**C**) Example AP trace illustrating waveform properties used for analysis. (**D**) Example voltage traces in response to hyperpolarizing current injections illustrating the quantification of input resistance and voltage sag ratio. (**E**) Principal component analysis (PCA) loading plot of demonstrating the relative contribution of each electrical property to each principal component. Membrane properties and spike kinetic parameters were used as variables for PCA. RMP, resting membrane potential. (**F**) Score plot of individual hDRG neurons, which clustered into three groups (E-types) based on their electrical properties. Each sample is colored by its E-type. (**G**) Pie chart depicting the percentage of hDRG neurons in each E-type cluster from a total of 228 recordings obtained from 56 independent donros. (**H**) Representative traces of AP waveforms from each E-type, shown in separate (left) and in overlay (right). (**I**) Representative phase-plane plots of the AP waveforms for each E-type. (**J-N**) Summary graphs of AP properties across the three E-types. N=33 (E-type 1), 74 (E-type 2), and 121 neurons (E-type 3). Bonferroni correction for multiple comparisons were used to determine threshold for statistical significance (α=0.005) for group comparisons using Kruskall-Wallis or ANOVA tests. P-values for Dunn’s post-hoc test are shown in plot. (**J**) AP half-width, p<0.0001 (significant), Kruskall-Wallis test. (**K**) Rise-time, p<0.0001 (significant), Kruskall-Wallis test. (**L**) AP decay time, p<0.0001 (significant), Kruskall-Wallis test. (**M**) Peak overshoot, p<0.0001 (significant), Kruskall-Wallis test. (**N**) AP rise-time, p<0.0001 (significant), Kruskall-Wallis test.

**Table 1.** Summary of hDRG donor demographic information.

| **N** | **56** |
| --- | --- |
| ***Race*** |  |
| **White** | **42** |
| **Black** | **12** |
| **Other/Unknown** | **2** |
| ***Sex*** |  |
| **Male** | **35** |
| **Female** | **21** |
| ***Cause of Death*** |  |
| **Anoxia** | **25** |
| **Stroke** | **19** |
| **Head Trauma** | **11** |
| **Other/Unknown** | **1** |
| **Age (years) †** | **45.63±15.28** |
| **Body mass index †** | **28.22±6.87** |

†Displayed in Mean ± SD

To classify the electrophysiological phenotypes (E-types) of hDRG neurons in an unbiased manner, we used principal component analysis (PCA) to cluster hDRG neurons based on their electrical properties (n=228; **Fig. 1E, F**). Features related to action potential (AP) kinetics, such as peak, decay time, and half-width, and features associated with intrinsic excitability, such as rheobase (the minimum current threshold required to fire an action potential), were found to have the largest contributions to the principal components (**Fig. 1E**). Using K-means neighbor clustering (**Fig. S1**), hDRG neurons were separated into three clusters: E-type 1, 2, and 3 (**Fig. 1F**). E-type 3 represented the largest cluster, made up of over half (121/228, or 53.07%) of the hDRG neurons patched. Approximately one-third (74/228, or 32.46%) of hDRG neurons were assigned to E-type 2, whereas E-type 1 consisted of a minority of hDRG neurons (33/228, or 14.47%; **Fig. 1G**). There were no significant associations between number of days in culture and the relative proportion of each E-type (**Fig. S2**). The membrane properties and action potential kinetics of the three E-types are summarized in **Table 2**.

**Table 2.**
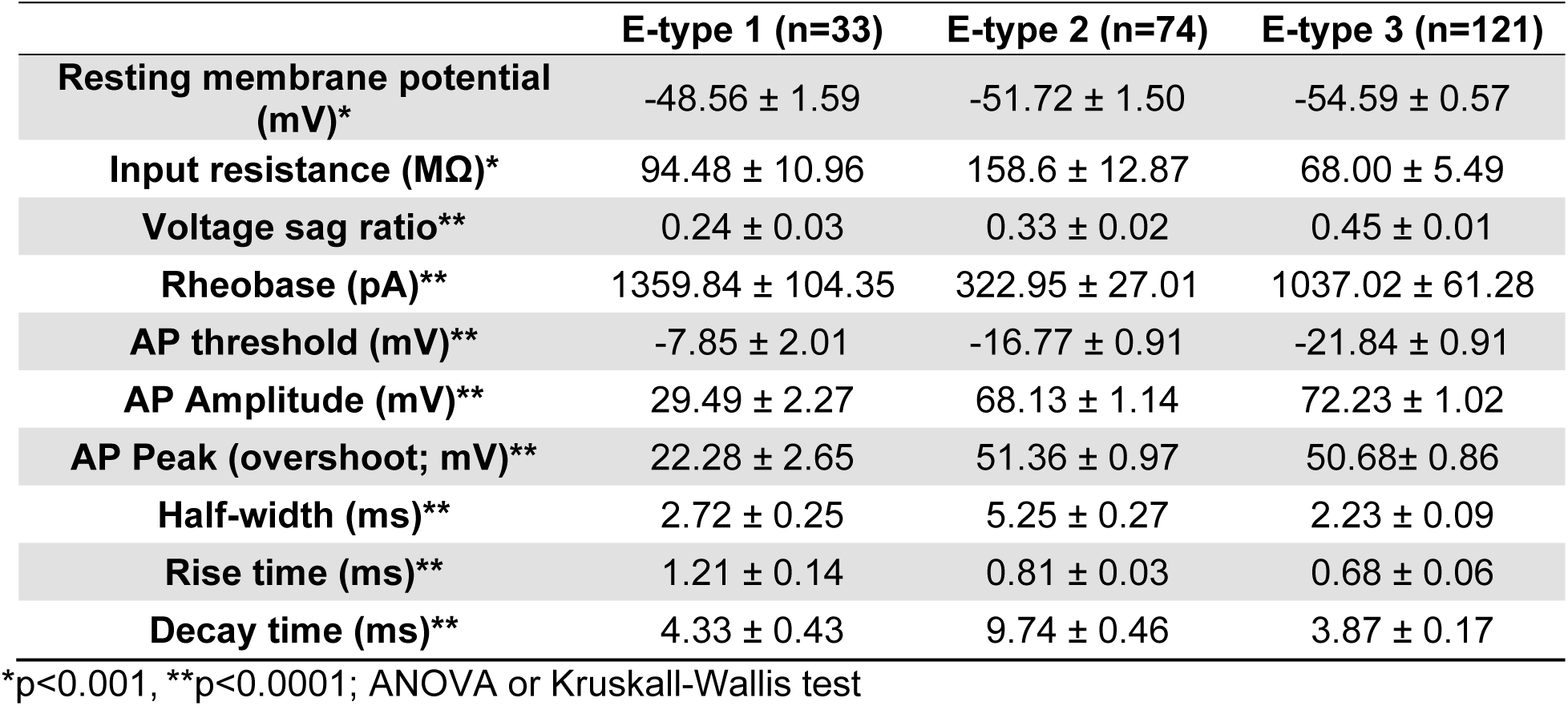
Summary of electrophysiological characteristics of each E-type.

Our PCA highlighted features associated with spike waveform and kinetics that contributed to clustering into three E-types (**Fig. 1E**). Analysis of the AP waveform kinetics (**Fig. 1C, H-N**) revealed that E-type 2 exhibits action potentials with significantly wider half-widths and a slower decay phase than both E-types 1 and 3 (**Fig. 1J, L**). While both E-types 1 and 3 were characterized by narrow action potential spikes with a fast decay phase (**Fig. 1H-J, K**), E-type 1 could be distinguished from E-type 3 based on its low peak and amplitude (**Fig. 1H, M, N**), as well as the slowest rise time of the three E-types (**Fig. 1I, K**). Together, these results suggest that hDRG neurons can be functionally clustered based on their membrane properties and spike kinetics.

### Human somatosensory neurons exhibit distinct high- and low-excitability E-types

Studies in both mouse and rat have reported at least 2-3 different firing subtypes within the small-diameter nociceptor population: one cluster that shows a non-repetitive, single-firing phenotype, and another that displays a repetitive firing pattern (*12*, *25*). These clusters may represent unique transcriptionally defined and functionally distinct subpopulations of somatosensory neurons (*25*). Additionally, these clusters have been hypothesized to represent different functional states of nociceptors, as the relative proportion of these clusters can change in response to nerve injury (*12*) or inflammatory mediators (*13*). Therefore, we analyzed if each hDRG E-type was associated with a distinct action potential firing pattern and excitability phenotype.

First, E-type 2 hDRG neurons fired action potentials in response to significantly lower current injections (**Fig. 2A-D**) and exhibited a significantly higher input resistance compared to E-types 1 and 3 (**Fig. S3A, B**), consistent with higher excitability. In contrast, hDRG neurons with E-type 3 had significantly higher rheobase than E-type 2 and exhibited the lowest input resistance, largest voltage sag, and the most hyperpolarized resting membrane potential out of the three (**Fig. S3B-D**), suggestive of low membrane excitability. Additionally, hDRG neurons with E-type 3 had a significantly larger cell diameter than neurons with E-type 2 (E-type 3 vs. E-type 2, 43.07µm ± 1.055 vs. 36.85 µm ± 0.97; **Fig. S4**).

**Fig. 2.**
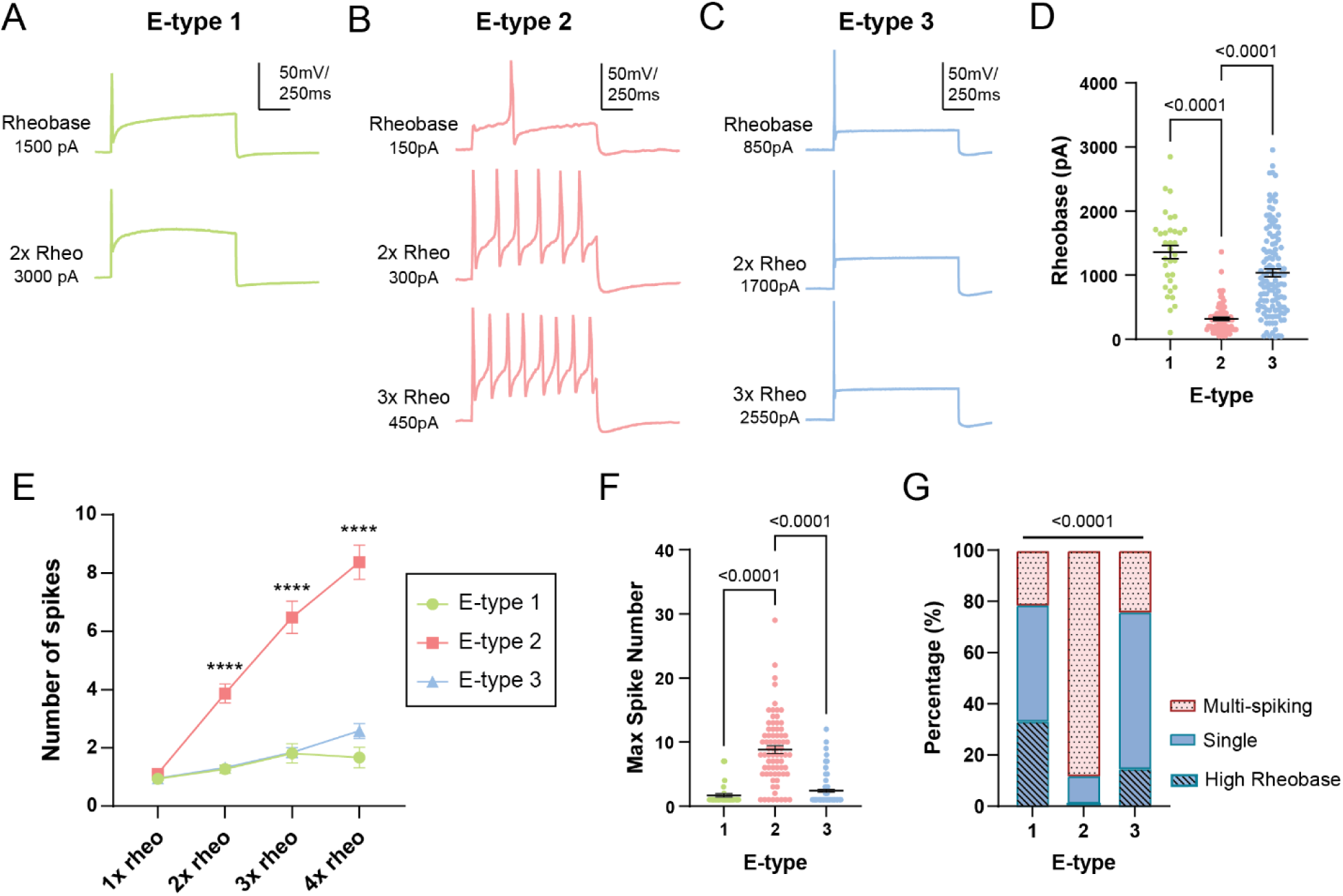
Action potential firing patterns in hDRG E-types. (**A-C**) Representative voltage traces of the AP waveforms in response to 1 second depolarizing current injections at threshold (rheobase) and suprathreshold (2-3x rheo) levels for each E-type. (**D**) Summary graph of rheobase values for each E-type. p<0.0001 (significant), Kruskall-Wallis test. P-values for Dunn’s post-hoc test are shown in plot. (**E**) Input-output curves of the number of AP spikes in response to 1 second current injections of 1-4x rheobase for each E-type). Linear mixed-effects analysis revealed significant fixed effect (p<0.0001; significant) of current injection, E-type, and current injection x E-type interaction. ****p<0.0001 (significant), Tukey’s multiple comparison test, E-type 1 vs. E-type 2, E-type 1 vs. E-type 3. (**F**) Summary graph of the maximum AP spike number in response to supratheshold depolarizing current injections for each E-type. p<0.0001 (significant), Kruskall-Wallis test. P-values for Dunn’s post-hoc test are shown in plot. (**G**) Percentage of the AP firing patterns (multi-spiking, single, or high rheobase) across each E-type. P<0.0001, Chi-square test. N=33 (E-type 1), 74 (E-type 2), and 121 neurons (E-type 3). Sample sizes are N=33 (E-type 1), 74 (E-type 2), and 121 neurons (E-type 3; **D-G**) Bonferroni correction for multiple comparisons were used to determine threshold for statistical significance (α=0.005) for statistical comparisons reported in (**D-F**).

Next, to determine each spike firing pattern, we analyzed the AP spike output to suprathreshold current injections (**Fig. 2E, F**). To determine the spike firing phenotypes across these three E-types, we classified neurons as multi-spiking (>1 action potential fired at up to 3x rheobase) or single-spiking (no more than 1 action potential fired at up to 3x rheobase). We observed that a subset of hDRG neurons had a significantly higher rheobase (1707 ± 90.9pA, **Fig. 2D, Table S2**) which limited us from testing their spike output at 3x rheobase; these neurons were classified as “high rheobase.” The relative proportions of multi-spiking, single-spiking, and high rheobase neurons were significantly different across the three E-types (**Fig. 2G**).

Analysis of spiking patterns revealed that E-type 2 fire a significantly higher number of action potentials at suprathreshold currents compared to E-types 1 and 3 (**Fig. 2A-C, E, F**), and E-type 2 neurons were predominantly composed of multi-spiking neurons (65/74, or 87.84%; **Fig. 2G**). In contrast, less than 25% of hDRG neurons with either E-type 1 (21.21%, or 7/33 neurons) or E-type 3 (23.97%, or 29/121 neurons) exhibited a multi-spiking pattern (**Fig. 2G**). E-type 3 neurons were most likely to be single-spikers (74/121, or 61.16%, **Fig. 2E-G**), whereas E-type 1 neurons were composed of both single-spiking (15/33 neurons, or 45.45%; **Fig. 2E-G**) and high rheobase neurons (11/33 neurons, or 33.33%; **Fig. 2G**). Together, these results suggest that E-type 2 represents a high-excitability, low-threshold, multi-spiking phenotype of human somatosensory neurons, whereas E-type 1 and 3 may represent populations of human sensory neurons with high activation thresholds.

### Patch-seq identifies four T-types of human DRG neurons

Recent scRNA-seq studies have revealed up to 22 transcriptionally distinct hDRG sensory neuronal subtypes, which can be broadly grouped into several major classes: A-fiber low threshold mechanoreceptors (A-LTMR), proprioceptors, A-fiber peptidergic neurons (A-PEP), C-fiber non-canonical peptidergic neurons (C-NP), and C-fiber peptidergic neurons (C-PEP) (*37*). However, it remains unknown whether distinct firing patterns are linked to specific transcriptionally defined populations (*1–4*). We thus performed Patch-seq recordings that allowed us to examine the relationship between E-types and transcriptional types or T-types from the same cultured hDRG cells (*39*). After patch-clamp recording, we performed RNA-seq on the cytosolic contents extracted from individually recorded cells (**Fig. 3A, Fig. S5A**). We established cDNA quality control (QC) metrics to identify high quality samples (**Methods, Fig. S5B**). Samples that passed the cDNA QC showed significantly more reads mapped to human genome and transcripts detected compared to those that failed QC (**Fig. S5C**). In total, we collected and processed 310 Patch-seq samples, and 139 passed QC metrics and thus were included in the transcriptional analysis.

**Fig. 3.**
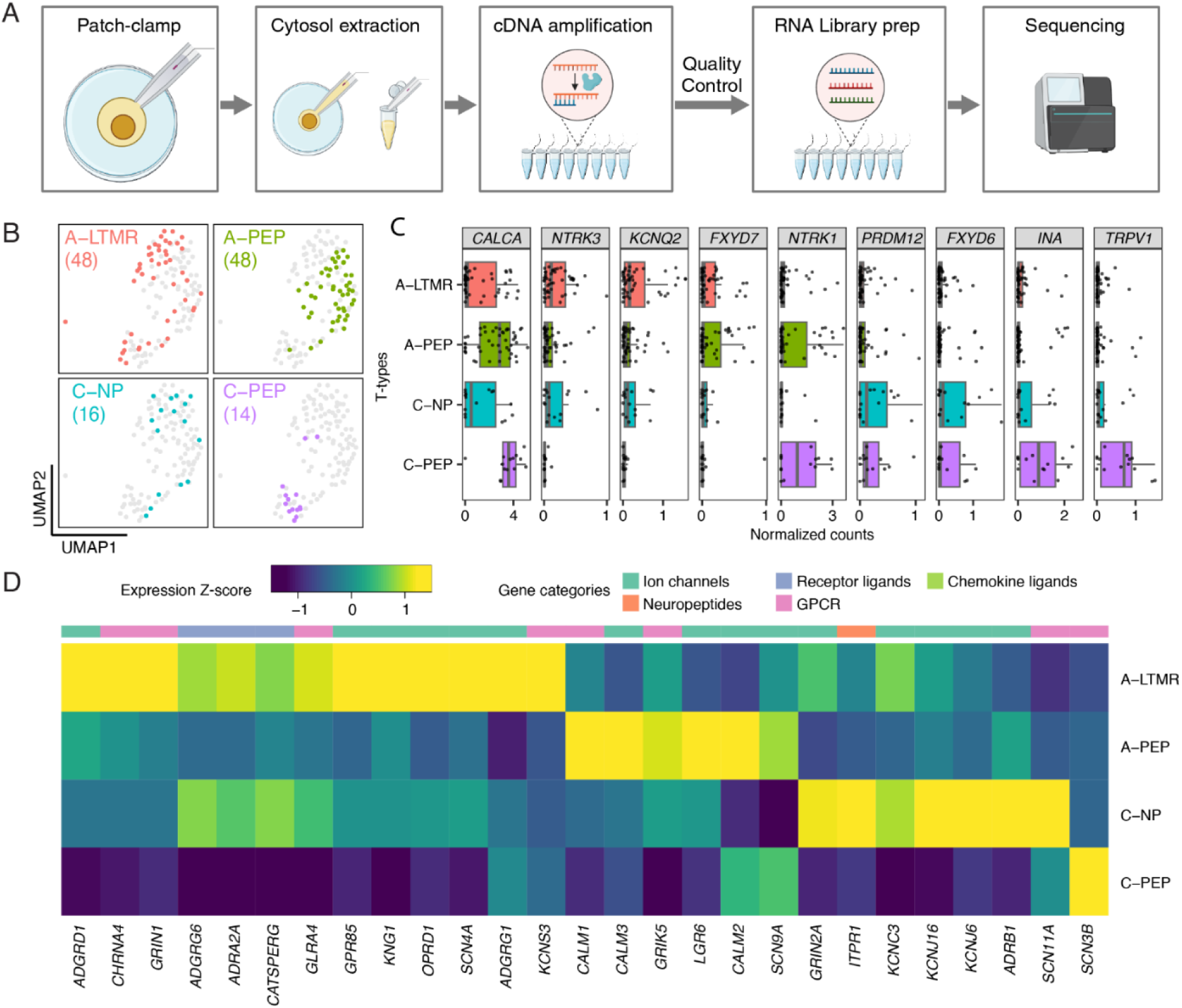
Patch-seq recordings identify four hDRG neuronal T-types. (**A**) Patch-seq workflow in human DRG neurons. (**B**) Uniform Manifold Approximation and Projection (UMAP) plot of 126 human DRG Patch-seq samples assigned to neuronal clusters. Samples are colored by T-types by anchoring and label transfer from the hDRG single-soma RNA-seq data, and the numbers in the parentheses indicate cell counts in each cluster. (**C**). Normalized counts (log2 transformed counts per million[log2CPM]) of select hDRG cell class marker genes in each T-type. Scatter plot indicates the expression in individual samples and the boxes indicate quartiles and whiskers are 1.5-times the interquartile range (Q1-Q3). The median is a grey line inside each box. (**D**) Heatmap showing the scaled expression (Z-score of log2CPM) of DEGs across T-types.

Despite recent efforts to elucidate the transcriptional profiles of diverse hDRG neuronal types, the evidence to provide the ground truth about their cytosolic RNA diversity is lacking as the majority of studies have focused on nuclear transcripts. We compared the average expression of all detected genes across Patch-seq samples to the average expression of those genes across neurons and non-neuronal cells from several hDRG single cell atlases previously reported, generated using varying input (e.g. single-nucleus, single-cell, and single-soma; **Fig. S5D**)(*28*, *32*, *34*, *36*, *41*). Consistent with their similarity in RNA origin, the transcriptional profile of the hDRG Patch-seq data best correlated to that of the hDRG single-soma RNA-seq data (*41*). Therefore, we used the cell type annotation and the marker genes identified from the single-soma RNA-seq data (*41*) to guide our transcriptional analysis.

To study the transcriptional identity of hDRG Patch-seq samples, we first performed clustering analysis that revealed four transcriptionally distinct clusters (**Fig. S6A**). Using the marker genes previously reported (*42*, *47*), we mapped each cluster to major human DRG cell classes (*36*, *42*). Cluster 1 to 3 express neuronal marker genes, such as *SNAP25* and *RBFOX3*, and are transcriptionally similar to hDRG neuronal cells, whereas cluster 4 expresses non-neuronal marker gene *APOE* and is transcriptionally more similar to non-neuronal cells previously described (**Fig. S6B, C**). Based on the expression of cell-type-specific DRG transcription factors and other canonical marker genes, clusters 2 and 3 likely correspond to A-PEP and C-PEP neurons, respectively, and cluster 1 appears to consist of transcriptionally distinct populations that match with *NTRK3*-expressing A-LTMR and *PRDM12*-expressing C-NP neurons (**Fig. S6D**). Notably, neuronal clusters also express injury-associated genes, such as *ATF3* and *SOX11*, which are likely induced by the cell dissociation and culture process (*43*). To better separate transcriptionally distinct hDRG neuronal classes, we next integrated the 126 Patch-seq samples from the neuronal clusters (clusters 1-3) to hDRG single-soma RNA-seq data and annotated their T-types by label transfer (**Fig. 3B**). T-type annotation largely corroborates the unbiased clustering (**Fig. S5**), and the transcriptional profiles of individual T-types are consistent with distinct hDRG neuronal classes previously reported (**Fig. 3C**). In addition, we identified 342 genes that are differentially expressed across T-types (**Table S3**), including 27 physiologically relevant genes (**Fig. 3D**).

### Patch-seq links E-type 2 to transcriptionally defined C-PEP hDRG neurons

Subpopulations of DRG sensory neurons exhibit significant differences in ion channel gene expression, modality-specificity, and conduction velocity(*14*, *16*, *32*, *34*, *44–46*). We next asked whether we can correlate hDRG electrophysiological and transcriptional identities. In total, 56 Patch-seq samples passed both electrophysiological and transcriptional analysis QC metrics and thus were assigned both E-types and T-types (**Fig. 4A**). These Patch-seq samples show similar E-type and T-type distributions compared to those of the larger populations examined solely in the electrophysiological or transcriptional analysis (**Fig. 4B,C**), despite the small sample size. When we examined individual T-types, the E-type distributions in A-LTMR, A-PEP, and C-NP neurons are similar to that of all Patch-seq samples when analyzed regardless of their T-types (**Fig. 4D**). Single-spiking, low amplitude E-type 3 is the most abundant in these T-types and constitutes more than half of the A-LTMR (61.9%) and A-PEP (58.3%) subpopulations. However, the C-PEP neurons had a significantly different E-type distribution from the other T-types, with most C-PEP neurons (83.3%) belonging to repetitive-spiking E-type 2, a 2.4-fold increase than other hDRG T-types. This suggests that C-PEP nociceptors are often low threshold, multi-spiking neurons. Consistently, E-type 2 Patch-seq samples exhibit significantly higher expression of hDRG nociceptor markers, such as *TAC1* and *CALCA* (**Fig. 4E, Table S4**), and its DE genes are important for the perception and response to pain (**Fig. 4F**). Taken together, our Patch-seq data suggests that hDRG C-PEP nociceptors are frequently multi-spiking E-type 2 neurons, whereas A-fiber neurons (A-LTMR and A-PEP) are more likely to exhibit high-threshold single-spiking phenotypes (E-types 1, 3).

**Fig. 4.**
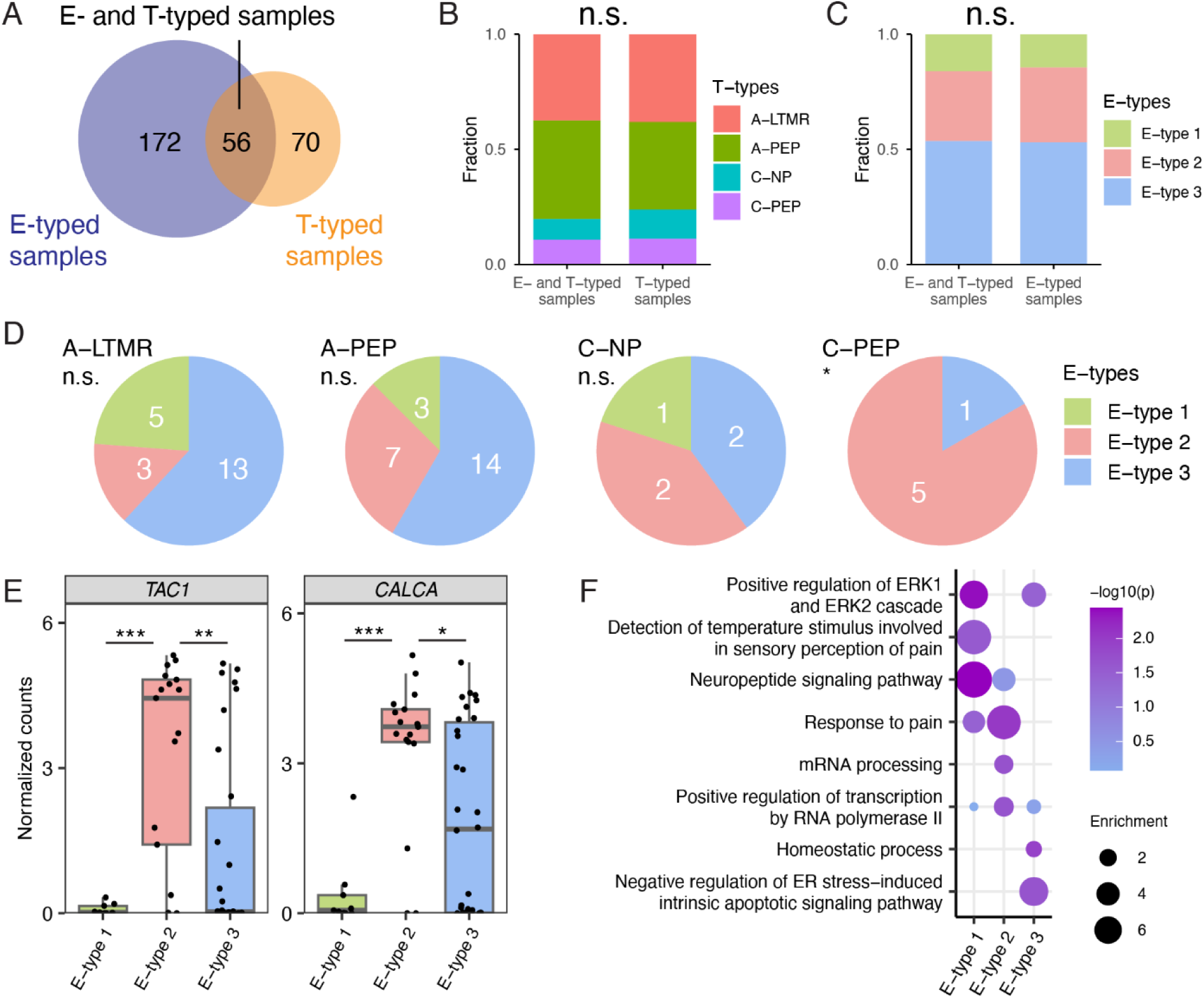
Patch-seq links hDRG E-types to T-types. (**A**) Venn diagram showing samples with E-types alone (included only in the electrophysiological analysis), samples with T-types alone (included only in the transcriptional analysis), and samples assigned both an E-type and T-type (included in both electrophysiological and transcriptional analyses). (**B**) Bar plot of the T-type distribution between samples assigned with T-types only and samples with both E- and T-types. Chi-square test, p = 0.8076. (**C**) Bar plot of E-type distribution between samples assigned with E-types only and samples with both E- and T-types. Chi-square test, p = 0.9129. (**D**) Pie chart showing the distribution of E-types in each T-type. Chi-square tests, p = 0.2415 for A-LTMR, p = 0.8593 for A-PEP, p = 0.8195 for C-NP, and p = 0.018 (*) for C-PEP. (**E**) Box plots showing the Normalized counts (log2 transformed counts per million[log2CPM]) of *TAC1* and *CALCA* across E-types. Scatter plots indicate the expression in individual samples and the boxes indicate quartiles and whiskers are 1.5-times the interquartile range (Q1-Q3). The median is a grey line inside each box. One-way ANOVA, p = 0.000212 for *TAC1* (***), and p = 0.000165 for *CALCA* (***). (**F**) Enriched gene ontology terms (p<0.05, enrichment > 1.5) across E-types.

### Donor pain history is associated with E-type-specific differences in action potential waveform and intrinsic membrane properties of human somatosensory neurons

Persistent nociceptor hyperexcitability in low threshold, multi-spiking DRG neurons is thought to drive persistent pain following nerve injury in rodents (*12*). Human somatosensory neurons resected from cancer patients with radiculo-neuropathic pain exhibit similar hyperexcitability and ectopic activity (*9–11*). However, it is unknown which cell types might underlie this ongoing pain, or whether these findings extend to other pain modalities like low back pain, which is the leading cause of disability worldwide (*47*). Therefore, we tested whether a donor’s history of pain was associated with any differences in membrane excitability and kinetics of repetitive and single-firing clusters of small- to medium-diameter hDRG neurons.

Donors were classified as having no significant pain history (“No Pain Hx”) or having a significant pain history in the dermatome of thoracolumbar DRGs (“Pain Hx”) based on the medical history reported by the next of kin (**Table 3, Table S1**). As the organ donors’ medical chart was not available through the organ registry paperwork, the medical history reported by the next of kin was used as a proxy. A donor was classified as having significant pain history if they met two out of three of the following criteria: (a) the word “pain” is used to describe donor’s recent or present medical condition (e.g., “daily pain,” “regular pain,” “painful tendonitis”), and the description of pain is aligned to thoracolumbar dermatomes (e.g., low back and/or the lower body); (b) donor’s medical history includes prescribed use of analgesics (e.g., hydrocodone or gabapentin) that is unrelated to substance use disorder, or there is documentation of other non-pharmaceutical interventions for pain (e.g., back surgery, nerve blocks); (c) donor’s medical history includes a disorder for which pain is a major symptom, such as arthritis or scleroderma. If the donor’s reported history was negative for all three criteria, the donor was classified as no significant pain history; donors who met one, but not all, of the criteria above were considered possible confounds and excluded from further analysis.

**Table 3.** Description of pain history in Pain Hx donors.

| Pain history description | Number of cells patched | Included in RNAScope |
| --- | --- | --- |
| Constant joint pain since age 17 | 0 | x |
| Leg pain and scleroderma, prescribed pain medication | 2 |  |
| History of multiple back surgeries, arthritis, bulging disc, back injury. Used prescribed pain medication for >5 years. | 3 |  |
| Knee pain due to arthritis | 1 |  |
| Knee pain due to tendonitis from playing football | 1 |  |
| Severe pain in the groin causing difficulty walking. Occasional migraines and pain in back of the neck. | 1 |  |
| Arthritis; history of knee surgery and prospective hip replacement surgery. | 2 | x |
| Degenerative disk disease; nerve block injections for back pain. | 8 | x |
| Arthritis; prescribed pain medication | 1 |  |
| knee pain, finger joint pain (unspecified laterality). "R hip problem" with neuropathy | 2 |  |
| Bulging lumbar disc managed with steroid injections in back and acetaminophen-hydrocodone | 1 |  |
| Fibromyalgia, osteoarthritis, rheumatoid/psoriatic arthritis with hx of R and L knee arthroscopy. Home medications include gabapentin, oxycodone HCL | 0 |  |
| Lower back pain managed by pain clinic, home medication includes Percocet | 16 |  |

Using the above criteria, we identified 13 donors (23.21%) as having a significant pain history (“Pain Hx”), which is close to the estimated prevalence of chronic pain among adults in the United States (*48*). Thirty-four donors were identified as having no significant pain history (“No Pain Hx”). Nine donors were excluded as possible confounds. The most common locations of pain were knee (5/13 donors) and low back (4/13 donors) pain (**Table 3**). The two most common etiology of pain were musculoskeletal conditions (5/13 donors; osteoarthritis, tendonitis, injury) and issues with intervertebral disks (3/13, “bulging” or “degenerative” disk; **Table 3).** There was no significant difference in the age, body mass index, or relative distributions of race, sex, and cause of death between the Pain Hx and No Pain Hx groups (**Table S5**). This suggests that low back pain and musculoskeletal pain are the most common chronic pain complaints seen in this organ donor group, which is consistent with population-level observations (*47*).

A total of 39 patch-clamp recordings in our E-type database were from hDRG neurons of Pain Hx donors. Upon post-hoc analysis, the majority of the recordings (61.54%) were obtained from two pain donors with low back pain (**Table 3**). Both groups had similar proportions of E-types 1, 2, and 3 (**Fig. S7**), suggesting that the overall distribution of the E-types does not vary between the two groups.

Next, we compared the electrophysiological properties of each E-type between the Pain Hx and No Pain Hx groups. Out of the 39 total patch-clamp recordings obtained from Pain Hx donors, only 4 neurons (10.53%) belonged to E-type 1 (**Fig. S7**), which represents a low-excitability phenotype that appears to map onto the A-LTMR and A-PEP T-types (**Fig. 1, 2**). The membrane properties, spike kinetics, and spike firing pattern of E-type 1 appeared to be largely similar across the two groups with the exception of rheobase, which trended higher in the E-type 1 neurons from the Pain Hx group (**Table S6, Fig. S8**). Statistical comparison was limited by the small sample size of E-type 1 in the Pain Hx group.

E-type 2 neurons exhibit a highly excitable multi-spiking phenotype (**Fig. 1, 2**) that strongly correlated with the C-PEP T-types and express high levels of human nociceptor markers (**Fig. 4**). The overall AP half-width, rise time, and decay time were not significantly different between the Pain Hx and No Pain Hx groups (**Fig. 5A-D**). However, we found that E-type 2 neurons from Pain Hx group had significantly larger action potential amplitude (**Fig. 5E, F**) and significantly higher voltage sag (**Fig. 5G, H**) compared to the No Pain Hx group. There were no significant differences in the overall spike firing, rheobase, input resistance, or resting membrane potential between the two groups (**Fig. 5I, J; Table S7**).

**Fig. 5.**
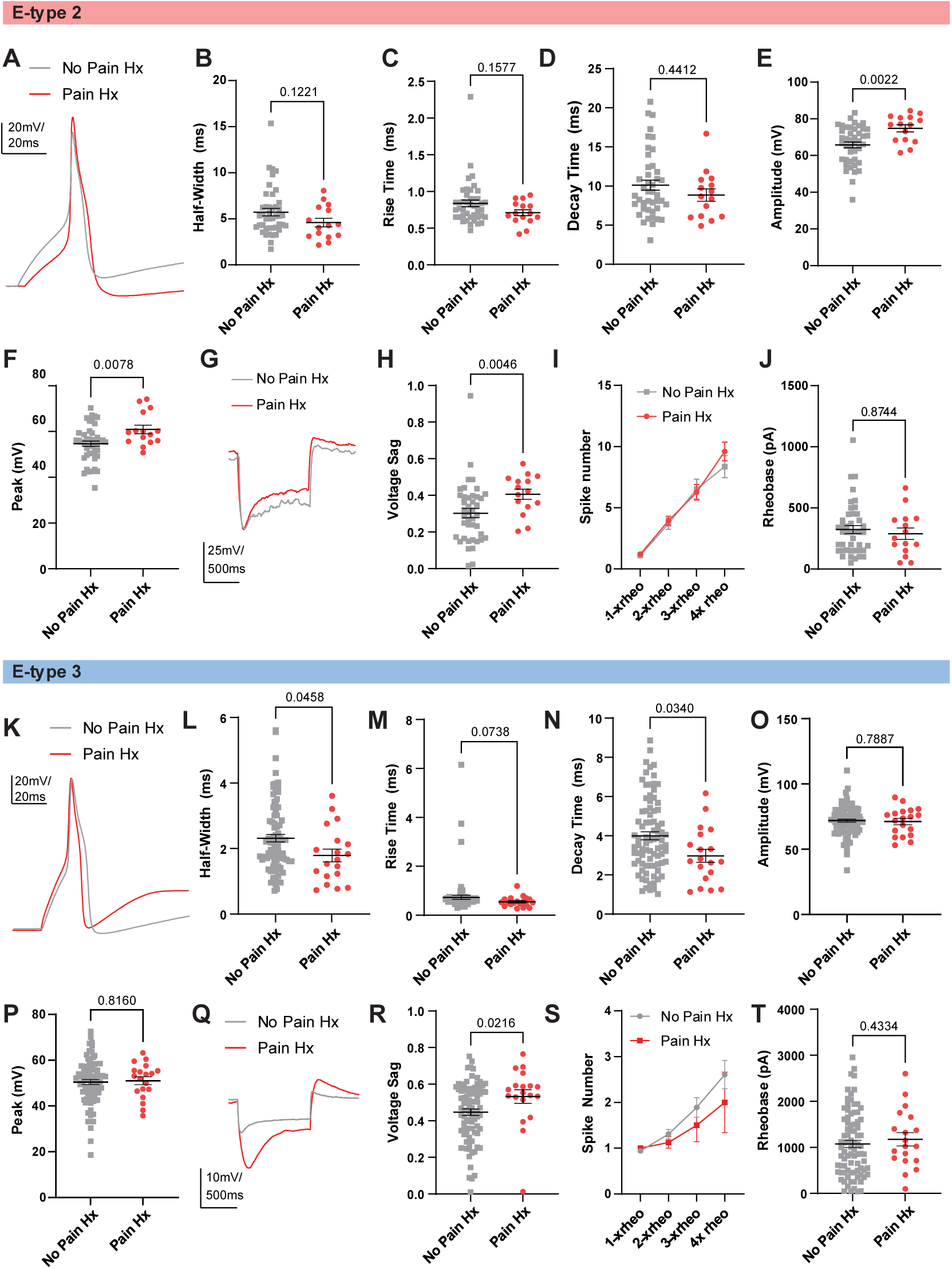
hDRG from donors with prior pain history exhibit E-type specific differences in their electrical properties. (**A-J**) Intrinsic membrane properties and AP kinetics of E-type 2 hDRG neurons, separated by donor history. N=42 (No Pain Hx), 15 (Pain hx). Bonferroni correction for multiple comparisons were used to determine the threshold for statistical significance (α=0.005) for group comparisons. (**A**) Representative AP traces elicited by 1 second depolarizing current injections at rheobase in hDRG E-type 2 neurons from donors with (red) and without (grey) prior pain history. (**B-F**) Summary graphs of AP kinetics of E-type 2 hDRG neurons, separated by donor pain history. (**B**) AP half-width; p=0.1221 (not significant), Mann-Whitney test. (**C**) AP rise time; p=0.1557 (not significant), Mann-Whitney test. (**D**) AP decay time; p=0.4412 (not significant), Mann-Whitney test. (**E**) AP voltage amplitude from threshold; p=0.0022 (significant), Mann-Whitney test. (**F**) AP Peak (overshoot); p=0.0078 (not significant), unpaired t-test. (**G**) Example voltage traces of E-type 2 neurons in response to hyperpolarzing current injections in hDRG from donors with (red) and without (grey) prior pain history. (**H-J**) Summary graphs showing voltage sag (H), input-output curves (I), and rheobase (J). (**H**) Voltage sag in response to hyperpolarizing current injections; p=0.0046 (significant), Mann-Whitney test. (**I**) Input-output curve showing number of spikes fired plotted against suprathreshold current injected. Mixed effects model analysis shows significant fixed effect of current injection (p<0.0001); fixed effects of pain history (p=0.5410) and the interaction of pain history x current injected (p=0.7277) were not significant. (**J**) Rheobase; p=0.4334 (not significant), Mann-Whitney test. (**K-T**) Intrinsic membrane properties and AP kinetics of E-type 3 hDRG neurons, separated by donor history. N=84 (No Pain Hx), 19 (Pain hx). Bonferroni correction for multiple comparisons were used to determine the threshold for statistical significance (α=0.005) for group comparisons. (**K**) Representative AP traces elicited by 1 second depolarizing current injections at rheobase in hDRG E-type 3 neurons from donors with (red) and without (grey) prior pain history. (**L-P**) Summary graphs of AP kinetics of E-type 3 hDRG neurons, separated by donor pain history. (**L**) AP half-width; p=0.0458 (not significant), Mann-Whitney test. (**M**) AP rise time; p=0.0738 (not significant), Mann-Whitney test. (**N**) AP decay time; p=0.0340 (not significant), Mann-Whitney test. (**O**) AP voltage amplitude from threshold; p=0.7887 (not significant), unpaired t-test. (**P**) AP Peak (overshoot); p=0.8160 (not significant), unpaired t-test. (**Q**) Example voltage traces of E-type 3 neurons in response to hyperpolarzing current injections in hDRG from donors with (red) and without (grey) prior pain history. (**R-T**) Summary graphs showing voltage sag (H), input-output curves (I), and rheobase (J). (**R**) Voltage sag in response to hyperpolarizing current injections; p=0.0216 (not significant), Mann-Whitney test. (**S**) Input-output curve showing number of spikes fired plotted against suprathreshold current injected. Mixed effects model analysis shows significant fixed effect of current injection (p=0.0021); fixed effects of pain history (p=0.2424) and the interaction of pain history x current injected (p=0.6400) were not significant. (**T**) Rheobase; p=0.4334 (not significant), Mann-Whitney test.

Finally, E-type 3 represents a cluster of hDRG neurons with low excitability and a characteristic narrow, fast action potentials that are biased towards A-PEP and A-LTMR T-types. E-type 3 neurons from Pain Hx group were associated with faster decay and a narrower half-width (**Fig. 5K-N; Table S8**); however, these differences did not meet the adjusted threshold for statistical significance after controlling for multiple comparisons (alpha=0.005, Bonferroni correction). Peak and amplitude were comparable between groups (**Fig. 5O, P; Table S8**). E-type 3 neurons in the Pain Hx group also showed trends towards a more depolarized resting membrane potential (**Table S8**) and a larger voltage sag (**Fig. 5Q,R**); however, other intrinsic membrane properties like rheobase, overall spike firing, action potential threshold, and input resistance were similar between the Pain Hx and No Pain Hx groups (**Fig. 5S,T; Table S8**). Together, these findings suggest that history of pain may be associated with E-type-specific differences in action potential kinetics in human somatosensory neurons, with the most pronounced differences seen in E-type 2, a cluster most strongly correlated with human nociceptors.

### Pain history is associated with higher NaV1.7 and 1.8 expression in human sensory neurons

Voltage-gated sodium channels NaV1.7, 1.8, and 1.9 are selectively expressed in the peripheral nervous system, act as key modulators of nociceptor excitability, and are thought to drive neuronal hypersensitivity that underlies persistent pain (*49*, *50*). The genes that encode these channels, *SCN9A* (encodes NaV1.7), *SCN10A* (NaV1.8), and *SCN11A* (NaV1.9), are selectively expressed amongst nociceptor subtypes in mouse, though they appear to be more broadly expressed in both nociceptor and non-nociceptor subpopulations in hDRG (*18*, *51–58*). As the expression of these voltage-gated sodium channels are upregulated in animal models of pain, (*59*) we investigated if there is pain history-associated differences in their expression in hDRG.

First, we conducted a preliminary analysis of the expression of *SCN9A*, *SCN10A*, and *SCN11A* using our Patch-seq dataset. After segmenting the dataset by donor’s pain history (**Table 3**), we observed trends towards higher *SCN10A* and *SCN11A* expression (16% and 33%, respectively) in hDRG neurons from donors with pain history (n=16; Pain Hx) compared to hDRG neurons from donors without pain history (n=73; No Pain Hx). However, given the small sample size, we were unable to further segment these results by E-types for differential expression (DE) analysis. Therefore, we opted to use a complementary approach to examining the distribution of *SCN9A*, *SCN10A*, and *SCN11A* expression in our donor population through RNA *in situ* hybridization. We quantified expression level in the DRG in two ways: (1) the overall percentage of DRG neurons that express the target gene, and (2) the relative expression level (e.g., number of mRNA copies of target gene) within each positive neuron. The Pain Hx group trended towards having a larger proportion of hDRG neurons expressing *SCN9A* and *SCN10A* compared to the No Pain Hx group, though the difference was not statistically significant (p=0.0655 and 0.0659, respectively; **Fig. 6A-B, E-F**). However, the expression levels of *SCN9A* and *SCN10A* per cell were significantly higher in the Pain Hx group compared to the No Pain Hx group (**Fig. 5C**, **G**). Increased copy numbers of *SCN9A* and *10A* were observed in small- and medium-sized hDRG neurons (**Fig. 5D, H**). While *SCN11A* expression was also higher in the Pain Hx group compared to the No Pain Hx group, as measured in terms of proportion and within-cell expression (**Fig. 5M-P**), this difference was not statistically significant. Interestingly, *SCN9A*, but not *SCN10A* or *SCN11A*, was detected in satellite glial cells surrounding neuronal cell bodies (**Fig. S9**), confirming previous observations (*11*, *42*).

**Fig. 6.**
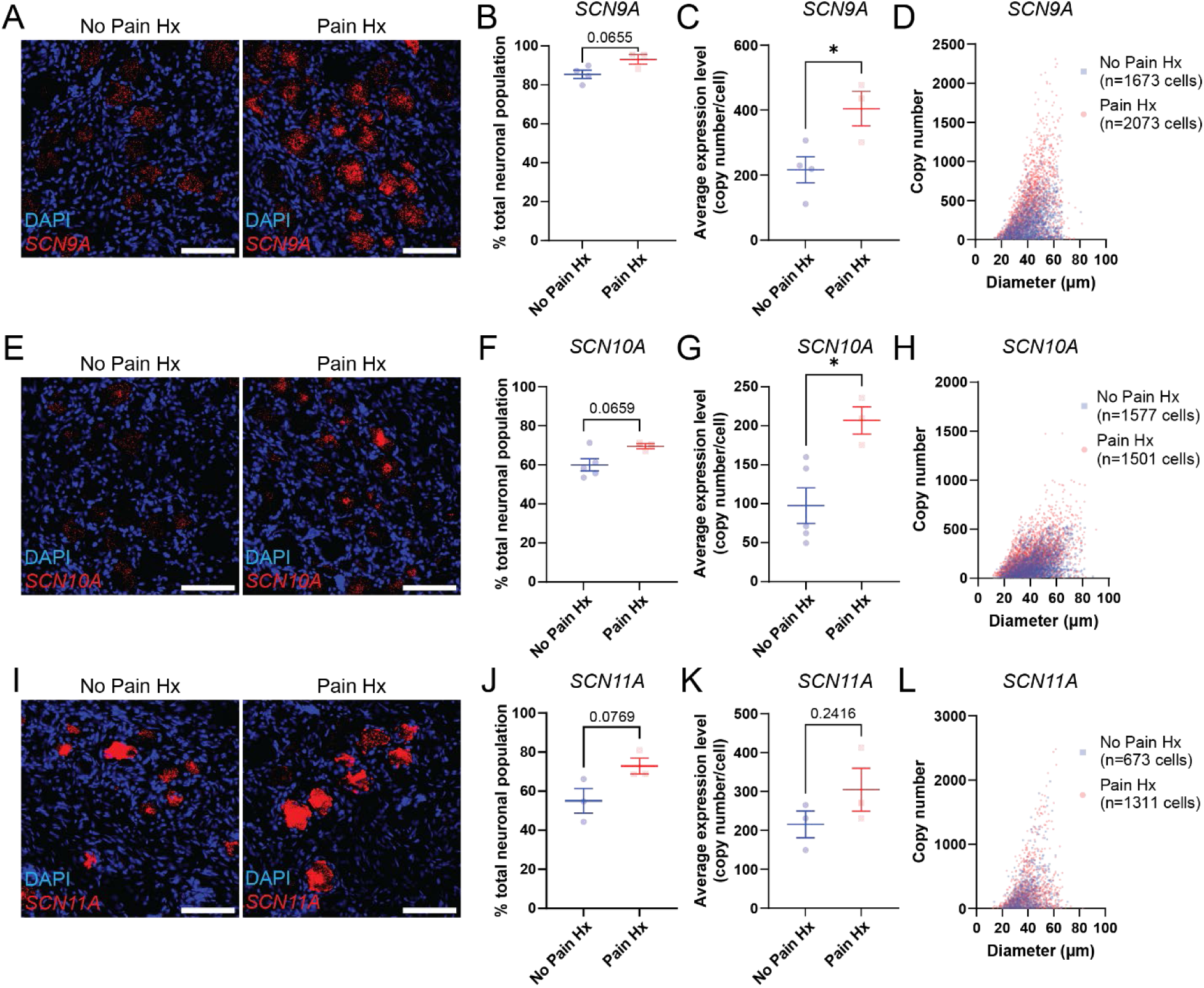
Donors with pain history exhibit higher levels of *SCN9A* and *SCN10A* expression in hDRG compared to donors without pain history. (**A**) Representative images of in situ hybridization for *SCN9A* (encoding Nav1.7, red) in hDRG sections from donors with (right) and without (left) significant history of pain. Scale bar = 100 µm. (**B**) Summary graph of the proportion of *SCN9A*+ hDRG neurons between donors with and without prior pain history. P=0.0655, unpaired t-test. N=4 (No Pain Hx), 3 (Pain Hx). (**C**) Summary graph of within-cell expression of *SCN9A* transcripts between donors with and without pain history. For each donor, the *SCN9A* copy numbers of all *SCN9A*-expressing neurons were averaged. N=4 (No Pain Hx), 3 (Pain Hx); 331-946 *SCN9A*+ cells per donor. (*) p=0.0343, unpaired t-test. (**D**) Scatter plot of *SCN9A* copy number vs. hDRG soma diameter for the two donor groups. N=1673(No Pain Hx), N=2073 (Pain Hx) neurons (**E**) Representative images of in situ hybridization for *SCN10A* (Nav1.8, red), in hDRG sections from donors with (right) and without (left) significant history of pain. Scale bar = 100µm. (**F**) Summary graph of the proportion of *SCN10A*+ hDRG neurons between donor groups. N=5 (No Pain Hx), 3 (Pain Hx); P=0.0659, unpaired t-test. (**G**) Summary graph of within-cell expression of *SCN10A* between donor groups. N=5 (No Pain Hx), 3 (Pain Hx); 134-627 *SCN10A*+ cells per donor. (*) p=0.0161, unpaired t-test. (**H**) Scatter plot of *SCN10A* copy number vs. soma diameter for the two donor groups. N=1557(No Pain Hx), N=1501 (Pain Hx) neurons. (**I**) Representative images of in situ hybridization for *SCN11A* (Nav1.9, red), in hDRG sections from donors with (right) and without (left) significant history of pain. Scale bar = 100µm. (**J**) Summary graph of the proportion of *SCN11A*+ hDRG neurons between donor groups. P=0.0769, unpaired t-test. N=3 for both Pain Hx and No Pain Hx groups. (**K**) Summary graph of within-cell expression of *SCN11A* between donor groups (3 donors per group, 175-719 *SCN11A*+ cells per donor). P=0.2416, unpaired t-test. N=3 for both Pain Hx and No Pain Hx groups. (**L**) Scatter plot of *SCN11A* copy number vs. soma diameter for the two donor groups. N=673 (No Pain Hx), 1311 (Pain Hx). List of Supplementary Materials

## Discussion

In the present study, we found that human sensory neurons exhibit three distinct E-types *in vitro*. Using Patch-seq, we mapped these electrophysiologically-defined E-types to transcriptionally-defined subpopulations of human sensory neurons, or T-types. Our findings indicate that the multi-spiking, high-excitability physiological subtype (E-type 2) is strongly correlated with C-fiber peptidergic (C-PEP) nociceptor subtype of hDRG neurons. Further, our investigation into the donor pain history revealed cluster-specific differences in action potential kinetics in hDRG neurons from donors with and without reported pain history, which may be partially explained by higher expression of *SCN9A* and *SCN10A* in lumbar hDRG of donors with pain history. Together, our work demonstrates how these types of multimodal datasets can be used to link functional phenotypes to different human sensory neuron populations and generate hypothesis-driven investigations into pain pathophysiology.

### Physiological heterogeneity of human sensory neurons

Multiple studies in both rodent and human DRG have classified nociceptors based on their electrophysiological features such as spiking patterns, action potential kinetics, presence of hump or shoulder in the rise or decay phase of the action potential, action potential firing latency, or presence of ectopic activity(*9*, *10*, *12*, *13*, *25*, *26*, *60*). In this study, we chose an unbiased clustering method that integrates multiple electrophysiological features. This approach avoids bias that may occur with subjective classification of E-types. This also allowed us to detect subtle differences in AP waveforms that might be influenced by distinct combinations of voltage-gated channels. Additionally, utilizing a large pool of donors for our sample collection allowed us to reduce potential donor bias in the classification process.

Echoing prior studies in rodents(*12*, *25*) and hDRG cultures (*61*), our analysis found that hDRG neurons demonstrate a wide variety of electrophysiological characteristics and evoked action potential firing patterns. In our pool of small diameter hDRG neurons (<60µm), three E-types emerged, with E-type 2 representing a highly excitable, multi-spiking subset a hDRG neurons and E-types 1 and 3 representing two clusters of low excitability neurons. Our control analyses suggest that distribution of E-types is not significantly related to number of days in culture seen under our experimental conditions, suggesting that number of days *in vitro* does not strongly influence E-type of each hDRG neuron between 2-11 days in culture.

It is unknown whether these E-types represent discrete, immutable physiological classes of hDRG neurons, or whether they reflect different possible electrophysiological *states*. Prior studies on spiking patterns provide some insight: for instance, persistent (i.e., 24hr) depolarization or prolonged exposure to inflammatory mediators can lead to changes in relative representation of single or repetitive-firing phenotypes in cultured human and rodent DRG neurons (*60*, *62*). This switch in firing patterns is not seen following acute exposure to inflammatory mediators (e.g., bradykinin) in hDRG neurons, which imply that spiking pattern changes may only occur in response to strong and persistent stimuli (*13*). While spike pattern is only one of many features that are incorporated in our E-type classification, these past studies suggest that E-types may reflect different functional states of hDRG neurons. This is further supported by our finding that T-types map onto more than one E-type.

The idea that E-types reflect functional states rather than a fixed phenotype may appear to be in conflict with our finding that E-type 3 neurons are significantly larger than E-type 2 neurons. However, our patch-seq analysis shows that A-LTMR and A-PEP, two T-types associated with large-diameter hDRG cells, are biased towards E-type 3. Then, it is possible that T-types are biased towards certain E-type at baseline, but that this E-type – or the electrophysiological state – may be plastic (*13*, *60*).

### Physiological specialization in transcriptionally distinct clusters of human sensory neurons

Linking the electrophysiological and transcriptional phenotypes in hDRG poses unique challenges due to the lack of readily available genetic labelling strategies, and the widespread distribution of nociceptors across multiple cell sizes. Patch-seq recordings allowed us to circumvent these challenges and obtain both electrophysiological and transcriptional data from the recorded hDRG neuron.

Recordings from genetically labeled mouse lines have revealed markedly different electrophysiological characteristics across DRG cell types (*25*). Each genetically labelled subpopulation of mouse sensory neurons is often associated with one distinct physiological phenotype, with the exception of peptidergic nociceptors which appear to be heterogeneous (*25*). In contrast to the earlier findings in mouse, we observed that hDRG T-types tend to map onto multiple E-types (**Fig. 4**), which suggests that transcriptionally defined subtypes of human sensory neurons are physiologically heterogeneous. Intriguingly, C-PEP – which reflects peptidergic nociceptors, a subpopulation which is thought to be more physiologically heterogeneous than other DRG classes (*25*, *26*) -- was the only T-type that had a significantly different E-type distribution and the only T-type to be predominantly made up of E-type 2, the highly excitable, multi-spiking E-type. Further studies are required to determine whether this is attributable to experimental conditions, such as differences in cell-type resolution between the studies or alterations in cell type marker gene expression *in vitro*, which may confound cell-type annotation in Patch-seq analysis (*63*). Alternatively, this departure from findings in mouse DRG may reflect the much more varied genetics, environments and histories (health status, exposures, injuries) of the human donors compared to inbred rodent lines raised in highly controlled environments. Finally, these differences may represent yet another cross-species difference in somatosensory neurons in addition to the transcriptional and molecular differences that have been characterized (*31–37*).

As the field continues to define the molecular features that distinguish hDRG cell types, it is possible that a given transcriptional class can include distinct subtypes based on their physiological or anatomical properties. While determining the axonal projections of defined DRG cell types will be a major hurdle in humans, information about soma diameter or axonal branching *in vitro* is readily available and may help to further distinguish specific cell types(*34*, *38*). In this study, we focused on smaller diameter neurons that are more likely to be nociceptors(*34*) and this approach can be expanded to study other cell types within the hDRG. In future studies, it will be valuable to incorporate different physiological stimuli, like temperature or pressure, to further characterize the response properties of specific populations of human sensory neurons. Continued investigation into the link between physiological characteristics and transcriptional features may, in the future, allow researchers to more reliably predict the transcriptional phenotype of a hDRG neuron based solely on physiological properties: for instance, neurons from different T-types that show similar firing patterns (i.e., E-type 2 neurons from A-PEP vs. C-PEP clusters) may have subtle electrophysiological differences that could be revealed with large collaborative efforts across the field. We suggest that multimodal approaches combining molecular profiling through single-cell sequencing or qPCR with recordings from human sensory neurons will be an essential step to fully understand the cell type diversity of human DRG. This may also present an opportunity to more directly relate new observations in human tissue to the decades of knowledge about the underlying neurobiology of pain and somatosensation developed using more tractable animal models.

Rapid advancements in high-throughput technologies to profile human tissue at cellular resolution have led to the identification of increasingly nuanced cell types. The biological ground truth of what defines a cell type is likely to require multimodal profiling across healthy and disease states. Previously, Patch-seq recordings have been used in rodent sensory neurons to identify mechanisms of mechanosensitivity (*40*) and human pluripotent stem cell-derived sensory neuron cultures (*64*). Developing a clearer picture of the features that define native human neurons will be useful to guide reprogramming efforts to better model and understand human disease using patient cells. Moreover, excitability changes within molecularly defined cell types could be used to identify different neuronal populations that could be drivers of specific chronic pain conditions, and potential therapeutic targets selectively expressed within those affected neurons. This will require harmonized datasets across large numbers of donors to fully realize this potential.

The data presented here represent an interim analysis of a planned larger effort to generate a large multimodal database of hDRG neurons that will eventually be used to compare sensory neuron characteristics within and across species. The dataset may provide additional predictive value to preclinical studies using cultured DRG neurons, as it may allow prediction of transcriptional/functional phenotype based on physiological markers. This type of detailed analysis of E-type relative to T-type represents a long-term goal of our NIH HEAL Initiative-supported human tissue program (part of the PRECISION Human Pain Network) and will require hundreds of recordings collected over several years to know whether specific electrophysiological features of hDRG neurons can be used exclusively to identify the recorded cell type. At present, we suggest that adding Patch-seq to recordings from human sensory neurons affords the opportunity to analyze physiological differences in genetically defined subsets of sensory neurons, reducing the impact of cell type heterogeneity and allowing for more direct comparisons of human sensory neuron biology to the cell-type-specific data collected from animal model studies.

### Pain-associated differences in neuronal excitability

To date, several electrophysiological analyses of dissociated hDRG neurons have been completed using samples obtained from cancer patients with neuropathic pain (*9–11*). Here, we aimed to use our dataset to see similar analyses could be conducted in samples obtained from organ donors.

Our study highlights both strengths and weaknesses of the approach of hDRG from organ donors for the study of somatosensation. In contrast to studies utilizing animal tissue, it is impossible to control for the entirety of a donor’s lived experience and there are likely many gaps in available medical histories. Therefore, it is possible that there could be chronic pain diagnoses or pain symptoms that were not reported. Our dataset almost certainly contains neurons that do not innervate the dermatome of pain in our pain donors. As a result, some of the heterogeneity seen in our Pain Donor dataset may be reflective of the heterogeneity in the innervation pattern of recorded neurons. Traditionally, innervation patterns of rodent DRG neurons have been identified through techniques such as genetic labeling and tracer injections which cannot be used in human subjects. However, as we develop increasingly more detailed understanding of hDRG gene expression patterns, we can utilize techniques like multiplexed immunohistochemistry techniques in various peripheral tissues such as skin and visceral organs to make predictions about the innervation patterns of hDRG subpopulations.=These are all inherent challenges in human-based research and will require the collective efforts of the pain research community to overcome.

Despite these limitations, the use of hDRG from organ donors allows us to more closely sample the chronic pain population at large. The prevalence of pain history in our donor pool was consistent with the estimated prevalence of chronic pain amongst adults in the U.S., and the pain history and location of chronic pain were consistent with the most common etiologies of chronic pain that is seen at a population level. For the present analysis, we included only donors with musculoskeletal pain conditions, but with continued efforts to build these data sets, eventually one can create more refined subcategories (e.g. only low back pain, which may have different drivers and involve different sensory neuron subtypes than arthritis, for example). While our analysis focused on pain history, our approach can be applied toward hypothesis-driven investigation based on other aspects of donor histories, such as substance use. One way to address the issue of heterogeneity is through a large sample size; a collaborative network of multiple labs and institutions with access to human tissue that can work together to contribute to a large, open database of hDRG neuronal recordings.

In our analysis, segmentation by pain history revealed E-type-specific differences in physiological features. The largest difference between groups was seen in E-type 2 neurons, which had significantly higher action potential peaks and larger amplitude. Action potential amplitude and peak are predominantly determined by voltage-gated sodium currents. In somatosensory neurons, NaV1.8 is the main determinant of the action potential amplitude: it contributes a large proportion of depolarizing currents during the rising phase of the action potential and has relatively depolarized steady-state inactivation, enabling robust spike amplitudes even at depolarized membrane potentials (*65*). Additionally, NaV1.7 may indirectly modulate the amplitude and peak by lowering the voltage threshold. The greater within-cell expression of *SCN9A* and *SCN10A* in the Pain Hx group suggest that upregulation of these two channels may underly some of the amplitude and peak differences seen in the Pain Hx group. Given that we did not see any differences in action potential threshold between the Pain Hx and No Pain Hx groups, NaV1.8 likely has a bigger contribution to the larger action potential height seen in the E-type 2 neurons of the Pain Hx group. In addition to the spike amplitude, Pain Hx was also associated with a larger voltage sag in E-type 2. While not statistically significant, E-type 3 neurons also showed similar trends towards larger voltage sag in the Pain Hx group. Voltage sag is primarily driven by hyperpolarization-activated cyclic nucleotide-gated (HCN) channels, which in turn can be modulated by cAMP signaling. Possible mechanisms that may underly the increased voltage sag seen in E-type 2 neurons from Pain hx donors include upregulation of HCN channel expression and potentiation of HCN-mediated current (I_h_) (*66–68*) due to altered GPCR tone. Since G protein pathways regulate numerous other ion channel and receptor families, including voltage-gated sodium channels, it will be important to directly compare expression and function between donor populations in future studies.

Our RNA *in situ* hybridization results point towards increased expression of NaV1.7 and 1.8 in hDRG obtained from pain donors. Unlike animal studies which enable greater experimental control which can provide more definitive conclusions about the directionality in the relationship between injury and ion channel expression, we are not able to draw causative conclusions due to the observational, rather than experimental, nature of our study involving donor-obtained hDRG neurons. However, a wealth of studies in animal models demonstrates that upregulation of NaV1.7 and 1.8 following injury or insult underlies the pathogenesis of persistent pain across multiple animal models (*11*, *69–72*). It is also possible that donors with Pain Hx had high levels of NaV1.7 and 1.8 at baseline, and that this predisposed them to developing chronic pain, as are seen in individuals born with gain-of-function mutations in NaV1.7 or 1.8 that predispose them to developing hereditary small-fiber neuropathies (*73–76*). The association between increased *SCN10A* expression and pain history is particularly intriguing considering the recent FDA-approval of the NaV1.8 inhibitor, suzetrigine (VX-548), which reduces post-surgical pain compared to placebo, with comparable efficacy to hydrocodone/acetaminophen (*77*, *78*).

In summary, our study identifies three distinct electrophysiological phenotypes of human somatosensory neurons, which can be linked to specific transcriptionally defined subpopulations using Patch-seq. Highly excitable E-type 2 appears to be strongly associated with the C-fiber peptidergic subpopulation of hDRG neurons. We additionally demonstrate that pain history is associated with E-type-specific differences in membrane properties and action potential amplitude, which may be due to higher levels of NaV1.7 and 1.8 expression. Detailed knowledge into how the properties of native human neurons are altered in chronic pain conditions may reveal new insights that can inform specific therapeutic strategies tailored to distinct etiologies of pain.

## Acknowledgements

We thank the past and present members of the Gereau lab for their helpful comments, and Mid-America Transplant for their collaboration in donor screening and tissue procurement. Finally, we thank the organ donors and their families, whose invaluable gift made this research possible.

This work was supported by the American Heart Association (AHA) through the AHA Predoctoral Fellowship #828671 (JY) and by the National Institutes of Health (NIH) through the NIH HEAL Initiative under award number U19NS130607 (RWG, BAC), part of the NIH PRECISION Human Pain Network. We thank the Genome Technology Access Center at McDonnell Genome Institute at Washington University School of Medicine for help with sequencing and data processing for the Patch-seq experiment. The Center is partially supported by NCI Cancer Center Support Grant #P30 CA91842 to the Siteman Cancer Center from the National Center for Research Resources (NCRR), a component of the National Institutes of Health (NIH), and NIH Roadmap for Medical Research.

## Author contributions

Conceptualization: JY, BAC, RWG

Data Curation: JY, LY, AC, KK

Formal Analysis: JY, LY, KK

Funding Acquisition: BAC, RWG

Investigation: JY, LY, AJW, AT, ZB, JDR, RAS, MP, AJD, JNL, RK, PG, JMM, GM, JL

Methodology: JY, LY, AJW, RWG, BAC

Visualization: JY, LY

Supervision: BAC, RWG

Writing—original draft: JY, LY

Writing—review & editing: JY, LY, BAC, RWG, AJW, ZB, JDR, RAS, AJD, JNL, RK, PG, AC

## Data availability

Donor metadata and electrophysiology data will be made available upon publication as an open repository via the SPARC Data Portal (doi.org/10.26275/thj5-bs3v) as approved by the NIH Data Coordination and Integration Center. Single-cell RNA sequencing data will be made available upon publication on dbGaP (accession number phs003788.v1.p1). Other data pertaining to this study will be made available upon reasonable request.

## Methods

### hDRG extraction

Extraction and collection of hDRGs were performed as previously described(*79*) in collaboration with Mid-American Transplant, with the following modifications and specifications. T11 - L5 DRGs were surgically removed from postmortem organ donors, within 1 −3 hours of aortic cross-clamping. Extracted DRGs were immediately placed in ice-cold, oxygenated N-methyl-D-glucamine (NMDG)-based artificial cerebrospinal fluid (aCSF; 93 mM NMDG, 2.5 mM KCl, 1.25 mM NaH_2_PO_4_, 30 mM NaHCO_3_, 20 mM HEPES, 25 mM glucose, 5 mM ascorbic acid, 2 mM thiourea, 3 mM Na^+^ pyruvate, 10 mM MgSO_4_, 0.5 mM CaCl_2_, 12 mM N-acetylcysteine; adjusted to pH 7.3 using NMDG or HCl, and 300 - 310mOsm using H_2_O or sucrose) and transported to lab for processing.

### Donor inclusion and classification

Donors that were older than 60 years, had a body mass index greater than 40, or had positive serology for blood-borne diseases such as hepatitis C or human immunodeficiency virus (HIV) were excluded from the study. Donor’s past medical history was determined using available information on the United Network of Organ Sharing (UNOS) documentation and interviews conducted by Mid-America Transplant with family members. A donor was classified as having significant pain history if they met two out of three of the following criteria: (a) the word “pain” is used to describe donor’s recent or present medical condition (e.g., “daily pain,” “regular pain,” “painful tendonitis,”), and the dermatome of pain is specified to be in the back and/or the lower body; (b) donor’s reported history includes prescribed use of analgesics (e.g., hydrocodone or gabapentin) that is unrelated to substance use disorder, or there is documentation of other non-pharmaceutical interventions for pain (e.g., back surgery, nerve blocks); (c) donor’s reportedhistory includes a disorder for which pain is a major symptom, such as arthritis or scleroderma. Based on these inclusion criteria, eight donors were identified as having a significant pain history (“Pain Hx”). Donors whose reported medical history did not mention any significant or possible pain conditions were classified as having no pain history (“No Pain Hx”). Finally, donors who only partially met the above criteria or had medical histories that included descriptions of possible pain-related conditions but lacked sufficient details to positively determine their pain statuses were considered to be possible confounds and excluded from further segmentation by pain history Demographic information and pain-related history of each donor is provided in **Table 1** and **Table S1**.

### hDRG primary culture

Primary culture of hDRGs were performed as previously described(*60*, *79*). Briefly, 2 - 3 thoracolumbar DRGs were cleaned and finely minced using ice-cold NMDG-aCSF. Minced DRGs were sequentially dissociated using papain (Worthington, Lakewood NJ) and collagenase type 2 (Sigma-Aldrich, Darmstadt, Germany), then triturated for mechanical dissociation. Dissociated DRGs were filtered and resuspended in DRG media (5% fetal bovine serum [Gibco, Grand Island, NY], 1% penicillin/streptomycin [Corning, Corning, NY], Glutamax [Life Technologies, Carlsbad, CA), and B27 [Gibco] in Neurobasal-A [Gibco]). DRG were plated and cultured on PDL and collagen-coated glass coverslips, and media was changed every 2 - 3 days.

### Patch-clamp electrophysiology

Patch-clamp experiments were performed between days in vitro (DIV) 2 - 11. Current clamp experiments were performed in external solution containing (in mM): 145 NaCl, 3 KCl, 2 CaCl_2_, 1 MgCl_2_, 7 glucose, 10 HEPES, adjusted to pH 7.3 with NaOH. Patch pipettes were pulled from thin-walled borosilicate glass with outer diameter of 1.5mm and internal diameter of 1.10mm (Sutter Instruments, Novato, CA). Pipettes were fire polished and had resistance values of 1 - 4 MΩ. The internal solution consisted of the following (in mM): 120 K-gluconate, 5 NaCl, 3 MgCl_2_, 0.1 CaCl_2_, 10 HEPES, 1.1 EGTA, 4 Na_2_ATP, 0.4 Na_2_GTP, 15 Na_2_Phosphocreatine; adjusted to pH 7.3 with KOH and HCl, and 290 mOsm with sucrose. Patch-seq experiments were conducted with a modified K-gluconate internal designed to promote cDNA yield while maintaining electrophysiological properties, containing the following: 110 K-gluconate, 4 NaCl, 10 HEPES, 0.2 EGTA, 1 Na_2_ATP, 0.3 Na_2_GTP, 10 Na_2_Phosphocreatine, 20 μg/mL glycogen, and 0.25 μL Protector RNAse Inhibitor; adjusted to pH 7.3 with KOH and HCl, and 290 mOsm with sucrose. Experiments were performed at room temperature. External solution was perfused continuously at a rate of 1 - 2 mL/min using a gravity-fed bath perfusion system. Recordings were made using a MultiClamp 700B amplifier, a Digidata 1550B digitizer, and the pClamp software (v.11.1, Molecular Devices, San Jose, CA). All recordings were sampled at 10 kHz.

hDRG neurons with soma diameter <60 µm were preferentially patched to increase the likelihood of recording from nociceptor populations(*34*). Membrane properties and excitability were measured in current clamp mode. After a stable whole-cell configuration was achieved and maintained for at least 2 minutes, a gap-free current-clamp recording was carried out for 1 - 2 minutes with zero current injection in order to assess resting membrane potential and spontaneous activity. After the resting membrane potential was measured, hDRG neurons were held at −60 mV for assessment of membrane excitability and passive properties. Membrane excitability was determined using 1-s long stepwise current injections starting at 50 pA (Δ 50 pA) until at least one action potential spike was elicited. To determine spiking pattern, suprathreshold currents at 1-, 2-, 3-, and 4-fold rheobase were injected using the same 1-s long stepwise protocol. hDRG neurons that had an unstable resting membrane potential, were depolarized >30mV at rest, or required more than −500pA to be held at −60mV were considered to be unhealthy and recordings from those cells were excluded from analysis for quality control.

Analysis of excitability, spike kinetics, and membrane properties was conducted using Easy Electrophysiology (Easy Electrophysiology Ltd., London, UK) and an in-house pipeline for automated analysis of electrophysiological data using the criteria described below. Rheobase was defined as the minimum current injection required to trigger an action potential firing. To assess spike firing pattern, the number of action potential spikes fired in response to each current step was counted to generate an input-output curve for each cell. The action potential threshold was defined as the membrane voltage when the first derivative of the voltage crossed 10 mV/ms (dV/dt = 10) and was determined from the first action potential fired at rheobase. Action potential kinetics were measured using the first action potential fired at rheobase. Maximum rise and decay times were obtained from 10 - 90% of the rising and falling phases, respectively. Action potential half-width was defined as the time between 50% of the rising and falling phases. Action potential amplitude was measured from action potential threshold to the peak of the spike, and peak was measured from 0mV to the peak of the spike. Input resistance was measured using hyperpolarizing current injections from −250 pA to −50 pA (Δ50 pA), by plotting ΔV_m_ against ΔI and determining the slope of the line of best fit. Voltage sag was measured using a single −250pA hyperpolarizing step. The magnitude of the sag was calculated as the difference between peak deflection and the steady-state V_m._ The sag ratio was calculated by dividing the sag by the peak deflection magnitude.

### Cytosol extraction for RNA-seq

After the conclusion of whole-cell recording, cytosolic contents were aspirated into the recording pipet using gentle negative pressure for 2 - 3 minutes, until a visible shrinkage of the cell was observed (Fig. S5A). After aspiration, the recording pipette was slowly and gradually withdrawn. Contents of the patch pipette were immediately ejected into a sample collection tube containing 1 µL 10 x Reaction Buffer (95% Lysis Buffer [Takara Bio] and 5% RNAse inhibitor [Takara Bio]) and 8µL RT-PCR grade nuclease-free water (Thermo Scientific) by applying positive pressure. Samples made of Patch-seq internal solution were used as negative controls to assess ambient RNA contamination. Samples were stored at −80 °C for long-term storage. For Patch-seq experiments, experimenters were agnostic to the pain history status of the donors.

### RNA sequencing and quality control of cytosolic contents

RNA was converted into cDNA using SMART-Seq v4 Ultra Low Input RNA Kit for Sequencing following the manufacturer’s manual. Samples containing 50 pg of total mouse brain RNA and samples made of nuclease-free water were used as positive and negative controls for library preparation, respectively. Twenty PCR cycles were used for cDNA amplification to increase yield. cDNAs were assayed using an Agilent TapeStation 4200. Samples were deemed high quality and proceed with sequencing library preparation if they met the following criteria: 1). containing no less than 1 ng of total cDNA, 2) at least 30% of total cDNA falling in the size range of 500 – 6000 bp, and 3). Both cDNA yield and cDNA percentage in range metrics were at least 25% higher than its associated negative controls.

The cDNA libraries that passed quality control were then prepared for sequencing using the Illumina Nextera XT Library Preparation Kit following the manufacturer’s manual. Resulting libraries were pooled and sequenced on an Illumina NovaSeq 6000 or NovaSeq X Plus instrument targeting at least 500k reads per library. Sequencing reads were aligned to hg38 Reference genome with STAR version 2.7.9a1. Gene counts were derived from the number of uniquely aligned unambiguous reads by Subread:featureCount version 2.0.32. Sequencing samples with at least 100k reads detected and at least 80% reads mapped to genome were included in the following analysis.

### Transcriptional clustering and T-type annotation of Patch-seq samples

In total, 139 samples that passed the cDNA and sequencing quality controls were included. First, the counts table of Patch-seq samples was loaded into Seurat (V4.4.0) in R (V4.4.1) for quality control and gene expression analysis (*80*). Raw counts were scaled to 10,000 transcripts per nucleus using NormalizeData() and scaled for each gene using ScaleData() functions. Highly variable genes were identified, and the top 20 principal components (PCs) were retrieved with RunPCA() using default parameters. Uniform Manifold Approximation and Projection (UMAP) coordinates were calculated using RunUMAP(). Clustering analysis was performed using FindClusters() based on the variable features from the top 10 principal components, with the resolution set at 1.2, and the marker genes for each cluster was identified using FindAllMarkers() comparing cells in one cluster to all other cells. Cell class markers were identified from single-soma RNA-seq data and used for annotating the transcriptional classes in Patch-seq data. Specifically, *SNAP25* and *APOE* were used to identify neuronal clusters and clusters with possible glial gene contamination, respectively. For neuronal classes, clusters with high expression level *CALCA* were identified as peptidergic neurons, and transcription factors, including *NTRK3*, *NTRK1*, and *PRDM12*, were used to identify A-LTMR, A-PEP/C-PEP, and C-NP/C-PEP neurons, respectively (*33*, *37*).

To increase cell type resolution, we directly compared the Patch-seq samples from neuronal clusters to single-soma RNA-seq data. First, FindTransferAnchors() was used to identify shared anchors (conserved features) between the datasets. T-type annotation was done using TransferData() that transferred the cell class labels (A-LTMR, A-PEP, C-NP, and C-PEP) described in the published dataset to each Patch-seq cell in the neuronal clusters.

### Differential expression (DE) analysis

DE analysis across T-types or E-types was done using DESeq2 wrapper function in Seurat comparing the gene expression of one group of cells to all other cells. Currently, the DE analysis is underpowered due to small sample size (126 sequenced samples, which were assigned to neuronal clusters were used for T-type DE analysis, samples with glial contamination were excluded, and 56 sequenced sample with E-type annotations for E-type DE analysis), and thus genes with average expression (Log2CPM) > 0.2 and adjusted p value < 0.1 were reported.

### Gene module scores

Gene module scores were calculated for each Patch-seq sample using AddModuleScore() in Seurat. To calculate the neuronal and non-neuronal module scores, DE analysis was performed comparing neuronal and non-neuronal cells from the hDRG cell atlases previously described (*37*). Top 200 marker genes based on Log2FC of neuronal and non-neuronal cells were selected. To calculate the injury scores, DE analysis was performed comparing the hATF3 cells to other cell types from the hDRG single-soma RNA-seq data previously described (*33*). Top 200 marker genes based on Log2FC were selected.

### Gene ontology (GO) analysis

GO analysis was performed using topGO (V2.40.0) in R. Differentially expressed genes described above were used as the input gene list and all genes with average expression >0.5 in the same cells are used as the background gene list. Genes were annotated for their biological process and GO terms with adjusted p value <0.05 were reported.

*In situ* hybridization

L1-L5 hDRGs were fixed in 4% paraformaldehyde at 4 °C overnight, cryopreserved in 30% sucrose, then flash-frozen in OCT. Fixed-frozen hDRG were cryosectioned at 12 μm and mounted on Superfrost Plus slides (Fisher, Pittsburgh, PA). Slides were stored at −20 °C or −80 °C with desiccant until further use. Immediately prior to beginning the *in situ* hybridization protocol (RNAScope, Bio-techne), hDRG sections were dehydrated using 50%, 70%, and 100% ethanol and treated with Protease IV and hydrogen peroxide (ACDBio, Minneapolis, MN). RNAscope was performed following manufacturer’s protocol. The following probes from ACDBio were used: SCN9A (cat. no.562251), SCN10A (cat. no.406291), and SCN11A (cat. no.404791). For each experiment, the probe was paired to Opal 570 (1:1000; Akoya Biosciences, Marlborough, MA). To reduce autofluorescence, TrueBlack Autofluorescence Quencher (Biotium, Fremont, CA) was applied at the end of RNAScope following manufacturer protocol, either before or after the application of DAPI. Slides were mounted with coverslips using ProLong Gold Antifade Mountant (Invitrogen, Waltham, MA) and stored at −20 °C until imaging.

### Imaging and image analysis

RNAScope slides were imaged using Leica DM6b system at 20 x magnification. For each slide, the entire hDRG section was imaged using the tile scan and merge feature of the using Leica Application Suite X (LAS-X, v.3.7, Leica Microsystems, Wetzlar, Germany). Acquisition settings were kept consistent across all slides and different experimental days. The following acquisition parameters were used: DAPI, 50 ms exposure, 2.1 gain, FIM 30%; L5, 300 ms exposure, 2.5 gain, FIM 55%; TXR, 500 ms exposure, 3.0 gain, FIM 55%. Segmentation and RNA expression analysis was completed using HALO AI^TM^ and the Multiple IHC module of HALO® (Indica labs). Satellite glial cells were distinguished from neuronal cell bodies visually based on their unique morphology.

### Statistics

All data were analyzed using GraphPad Prism 10 (GraphPad Software Inc, Boston, MA) and R (V4.4.1). To inform the use of appropriate statistical tests, normality of residuals and heteroscedasticity were assessed for each dataset using Shapiro-Wilk test and Brown-Forsythe test, respectively. To assess statistical differences across multiple groups, one-way ANOVAs with Tukey’s post-hoc test, Kruskall-Willis test with Dunn’s post-hoc test, or Brown-Forsythe ANOVA with Dunnett’s T3 tests were performed. For comparison of multiple electrophysiological features within the same set of samples, Bonferroni correction was applied to determine a multiple comparison-adjusted threshold (α=0.005) for statistical significance. All post-hoc tests were performed with multiple comparison corrections. To assess differences between groups in the input-output curves, restricted maximum likelihood (REML) mixed effects analysis with Geisser-Greenhouse correction was used with Tukey’s or Šidák’s multiple comparison tests for post-hoc analysis where appropriate. PCA analysis and K-means clustering of electrophysiological data was conducted using R. All data are represented as mean ± SEM unless otherwise noted. Detailed statistics for all comparisons presented in the study can be found in **Supplementary Table S9**.

### Study approval

Extraction procedures were approved by Mid-America Transplant, and an Internal Review Board (IRB) waiver was obtained from the Human Research Protection Office at Washington University in St. Louis. hDRG samples were obtained from postmortem organ donors with the consent for tissue donation for research from family members.

## List of Supplementary Materials

### Supplementary Tables

Table S1. Detailed donor information (see separate Excel file)

**Table S2.**
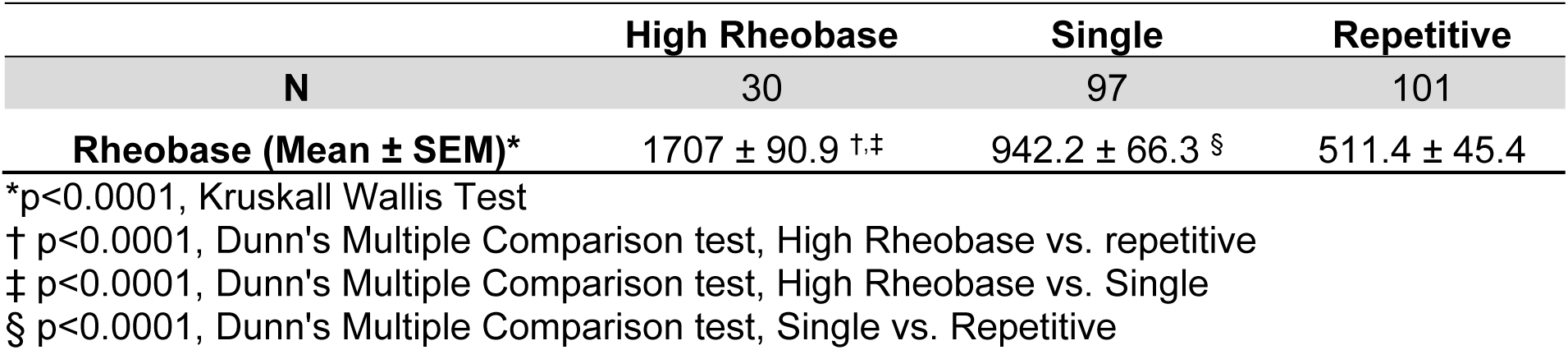
Comparison of Rheobase between hDRG neurons with different firing patterns.

|  | High Rheobase | Single | Repetitive |
| --- | --- | --- | --- |
| N | 30 | 97 | 101 |
| Rheobase (Mean ± SEM)* | 1707 ± 90.9 <sup>†,‡</sup> | 942.2 ± 66.3 <sup>§</sup> | 511.4 ± 45.4 |

\*p<0.0001, Kruskal Wallis Test
† p<0.0001, Dunn's Multiple Comparison test, High Rheobase vs. repetitive
‡ p<0.0001, Dunn's Multiple Comparison test, High Rheobase vs. Single
§ p<0.0001, Dunn's Multiple Comparison test, Single vs. Repetitive

Table S3. DEG T-types (see separate Excel file)

Table S4. DEG E-types (see separate Excel file)

**Table S5.**
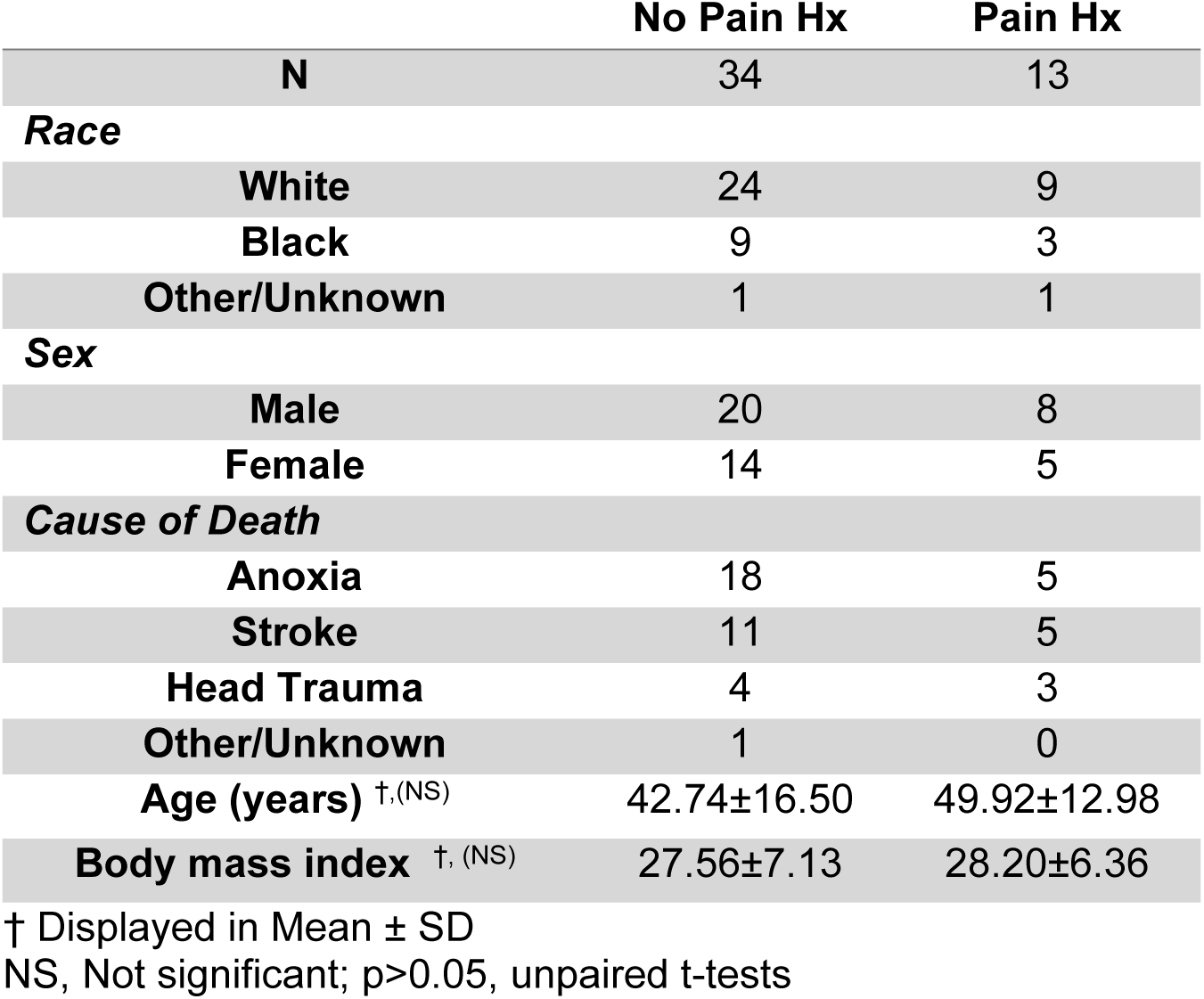
Summary of hDRG donor characteristics, segmented by pain history.

**Table S6.**
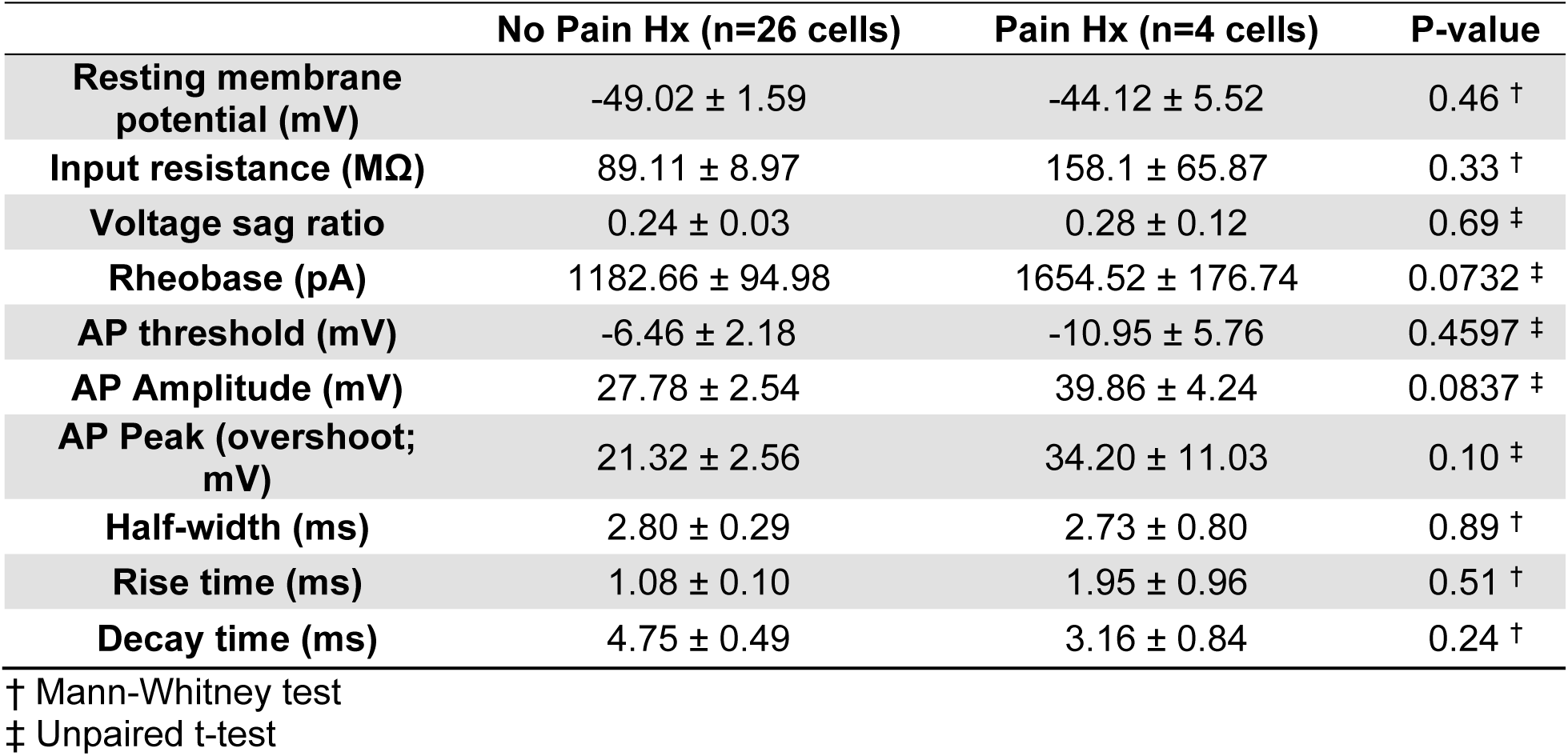
Summary of electrophysiological features of E-type 1 between Pain Hx and No Pain Hx donors.

**Table S7.**
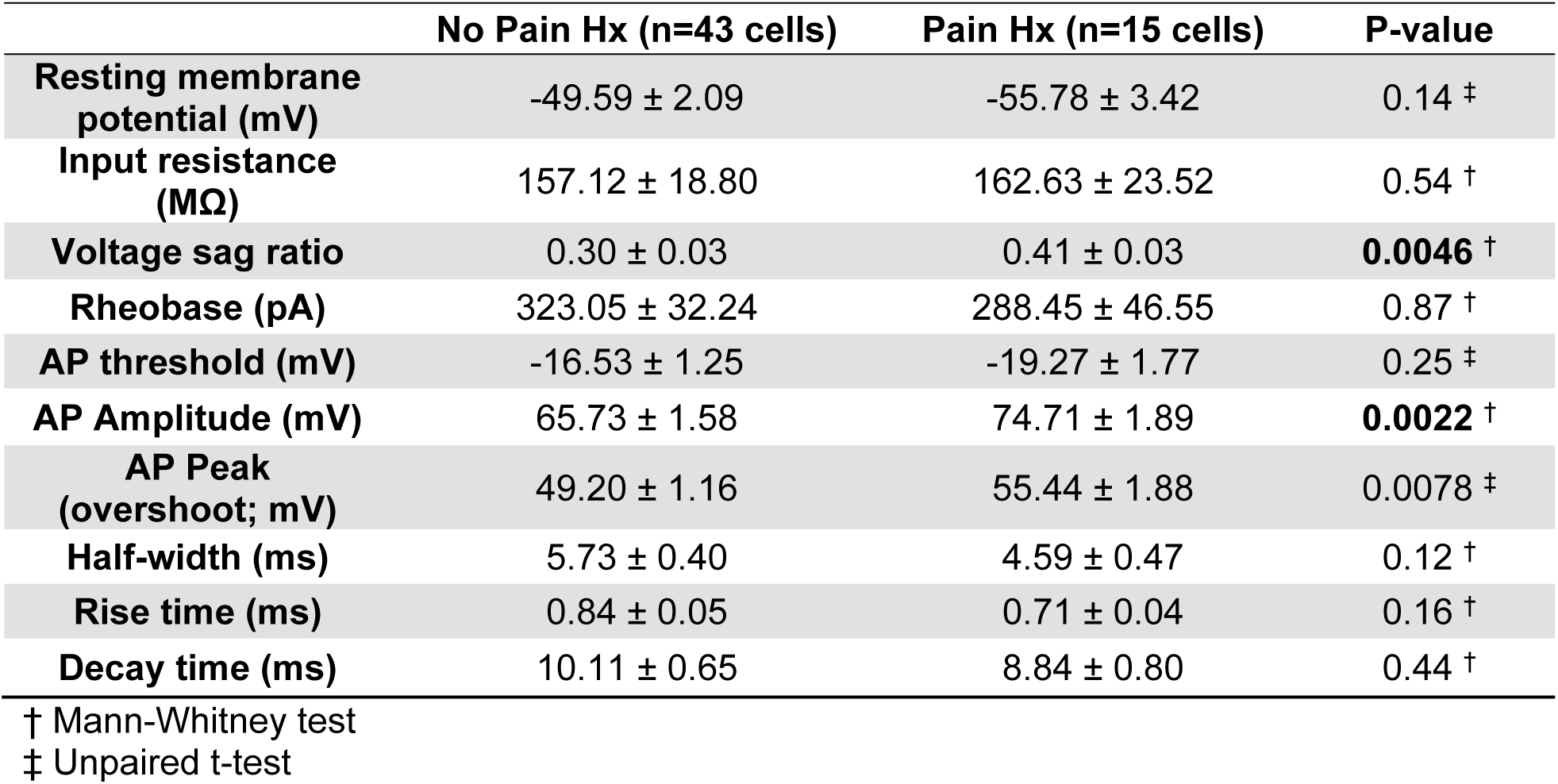
Summary of electrophysiological features of E-type 2 between Pain Hx and No
Pain Hx donors.

**Table S8.**
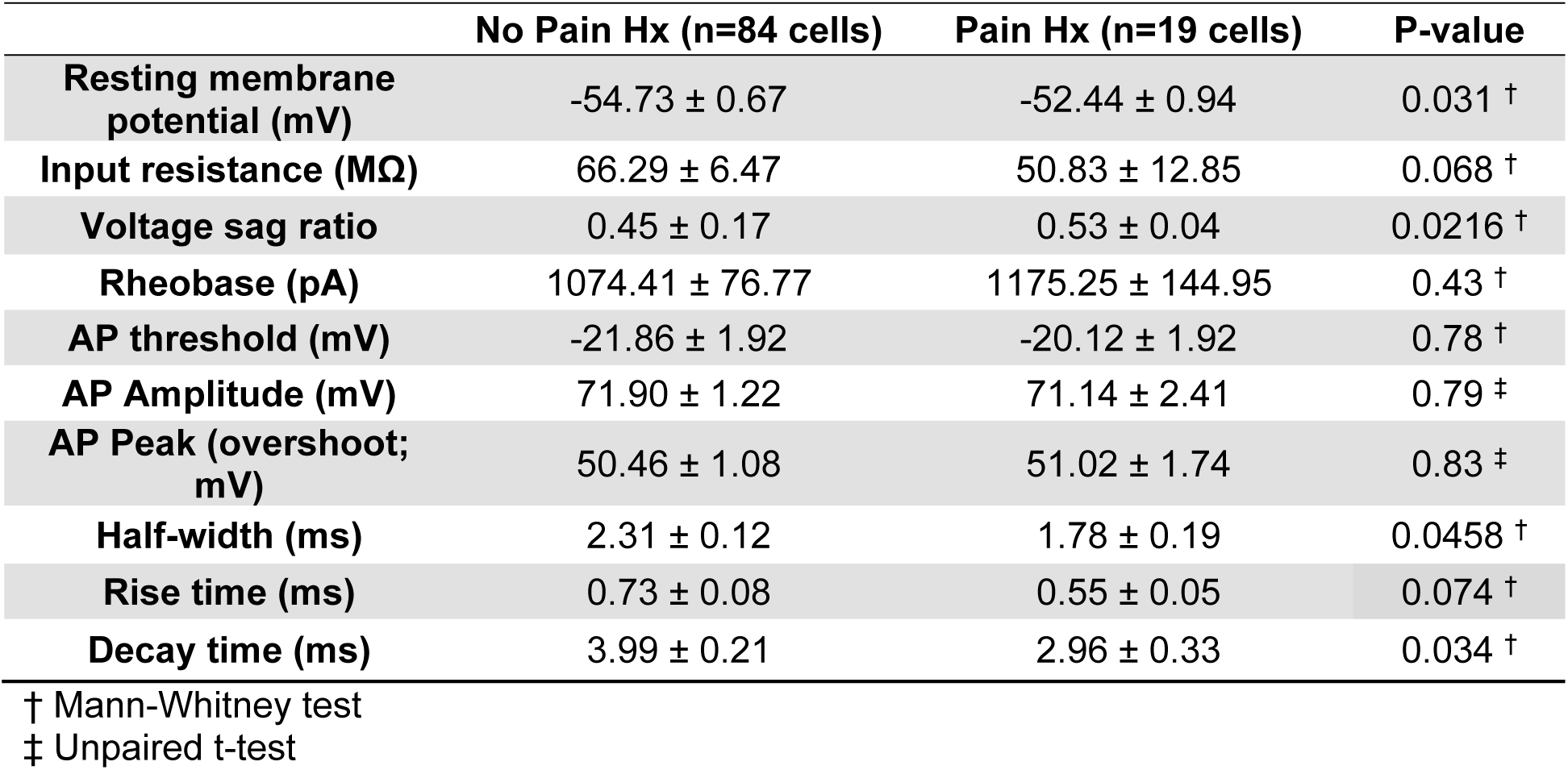
Summary of electrophysiological features of E-type32 between Pain Hx and No Pain Hx donors.

Table S9. Statistics Table (see separate Excel file)

### Supplementary Figures

**Fig S1.**
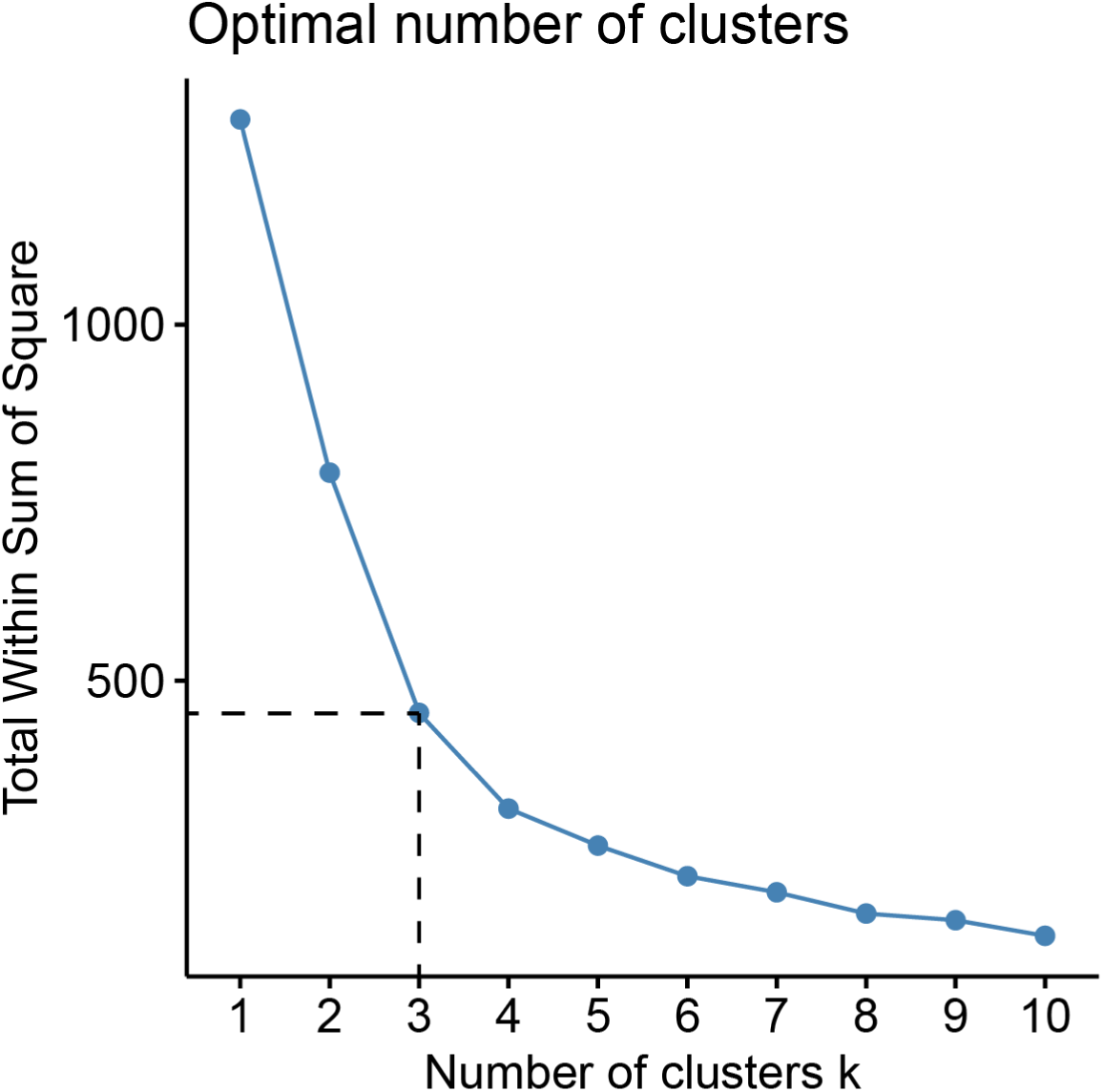
Elbow plot showing the within-cluster sum of squares against different K values. We chose k=3 as the ideal cluster number for balancing model fit and complexity.

**Fig S2.**
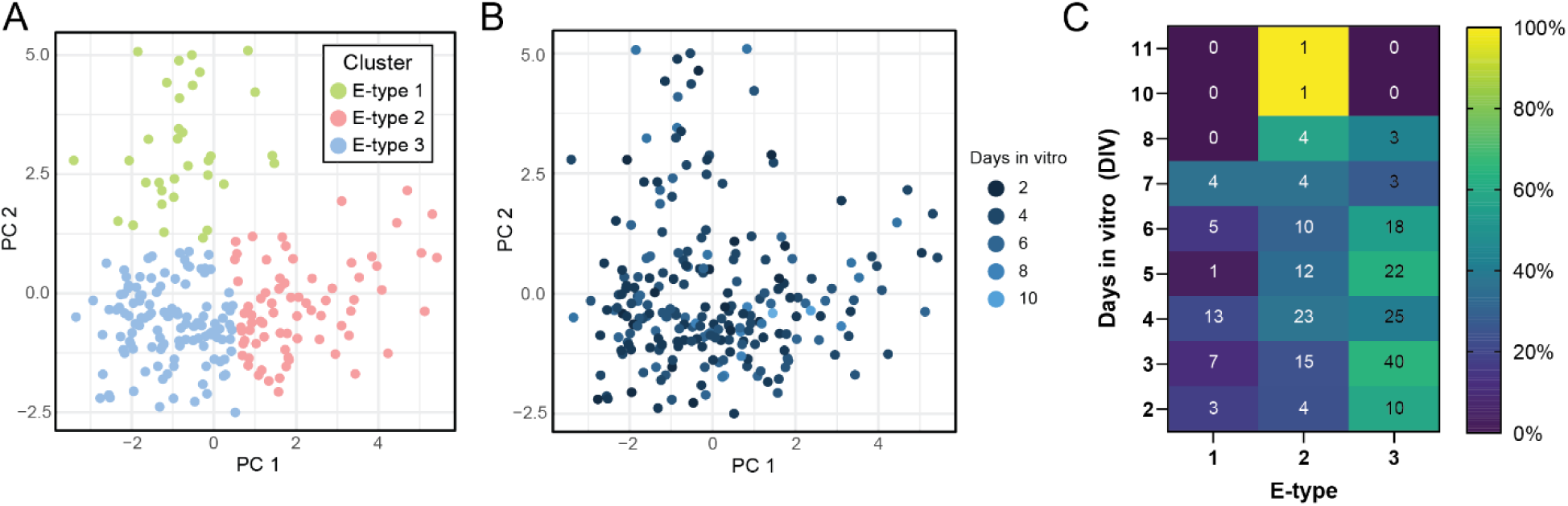
E-type distribution across days in vitro. (**A**) Score plot of individual hDRG neurons clustered by their electrophysiological properties. (**B**) Same plot in panel A, but each neuron is color-coded by the number of days in culture before recording. (**C**) Heat map of E-type distributions across DIV. Number in each box shows number of cells corresponding to the E-type on the X-axis that were recorded at the DIV specified on the Y-axis. Each box is color-coded based on the relative proportion (%) of each E-type out of all the cells recorded on a particular DIV.

**Fig S3.**
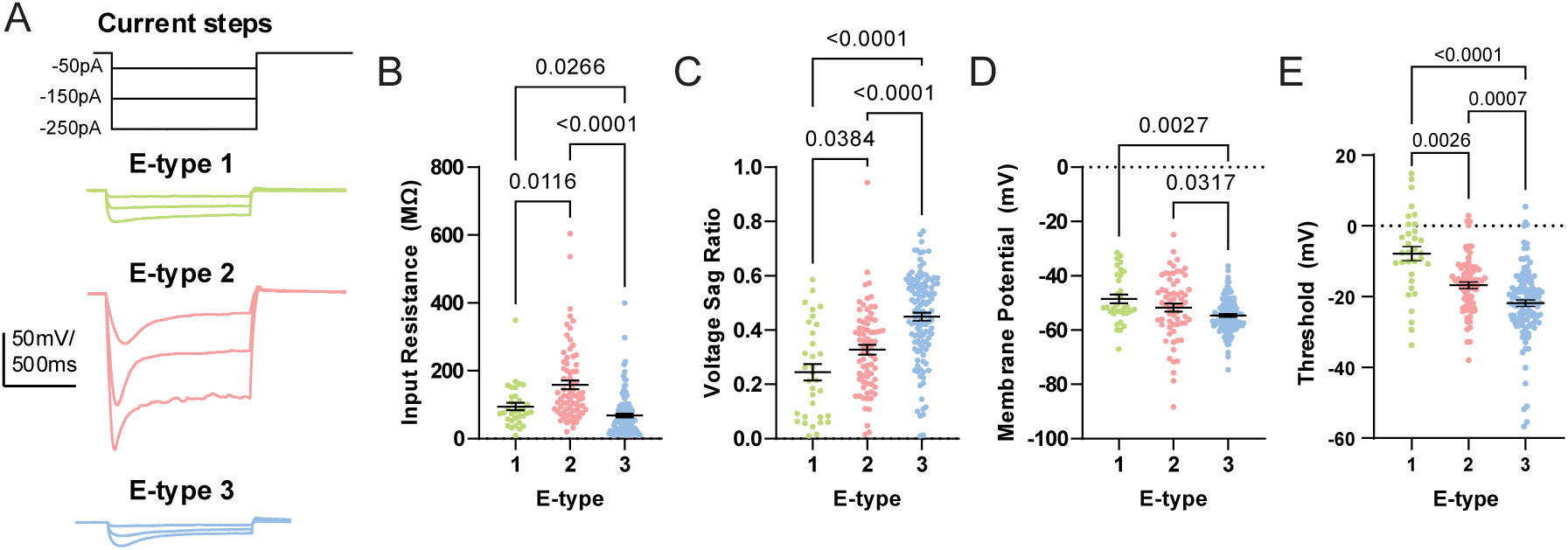
Additional membrane properties of hDRG E-types. (**A**) Example voltage traces of each hDRG E-type in response to 1 second hyperpolarizing current injection steps. (**B-E**) Summary graphs of input resistance (**B**), voltage sag (**C**), resting membrane potential (**D**), and AP threshold (**E**) for each hDRG E-type. N=33 (E-type 1), 74 (E-type 2), and 121 neurons (E-type 3). Bonferroni correction for multiple comparisons were used to determine threshold for statistical significance (α=0.005) for group comparisons using Kruskall-Wallis or ANOVA tests. P-values for Dunn’s post-hoc test (Kruskall-Wallis) or Tukey’s post-hoc test (ANOVA) are shown in plot. (**B**) Input resistance, p<0.0001 (significant), Kruskall-Wallis test. (**C**) Voltage sag ratio,, p<0.0001 (significant), one-way ANOVA. (**D**) Resting membrane potential, p<0.0001 (significant), Kruskall-Wallis test. (**E**) AP threshold, p<0.0001 (significant), Kruskall-Wallis test.

**Fig S4.**
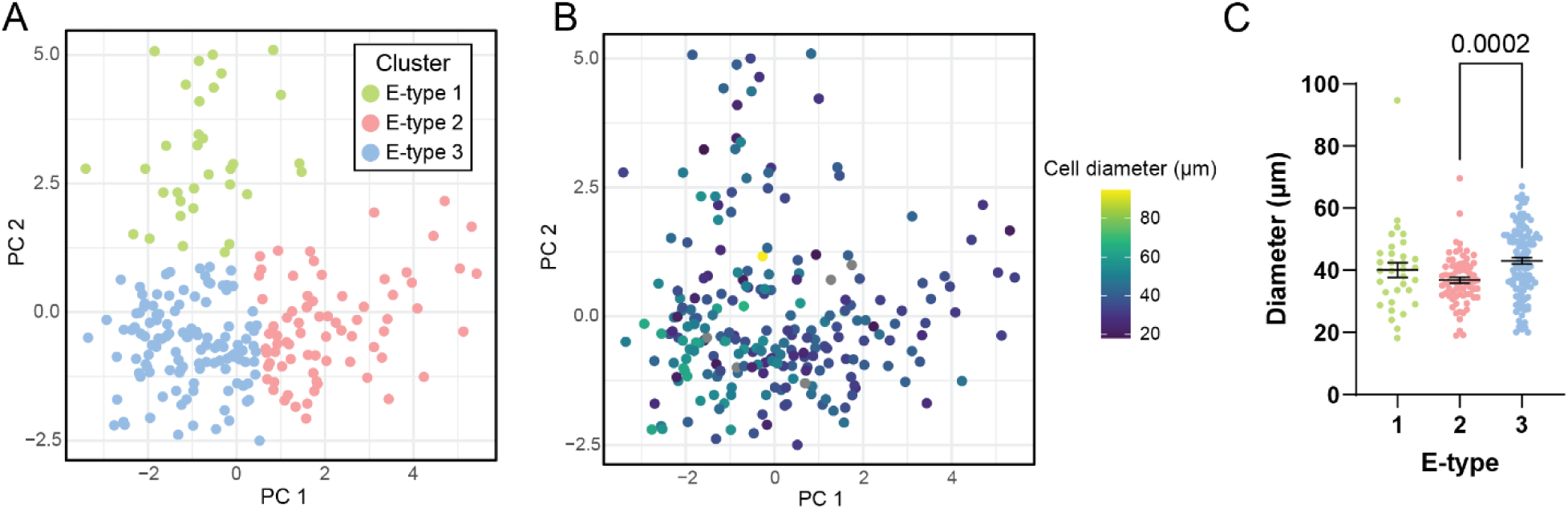
Soma diameters across hDRG E-types. (**A**) Score plot of individual hDRG neurons clustered by their electrophysiological properties. (**B**) Same PCA plot in (**A**) color-coded by a heat map of soma diameters in microns. (**C**) Summary graph of soma diameter for each hDRG E-type. P=0.0003, Kruskall-Wallis test. N=33 (E-type 1), 74 (E-type 2), and 119 neurons (E-type 3). Post-hoc comparisons: p=0.0002, E-type 2 vs. E-type 3; p=0.6449, E-type 1 vs. E-type 2; p=0.2671, E-type 1 vs. E-type 3; Dunn’s post-hoc test.

**Fig. S5.**
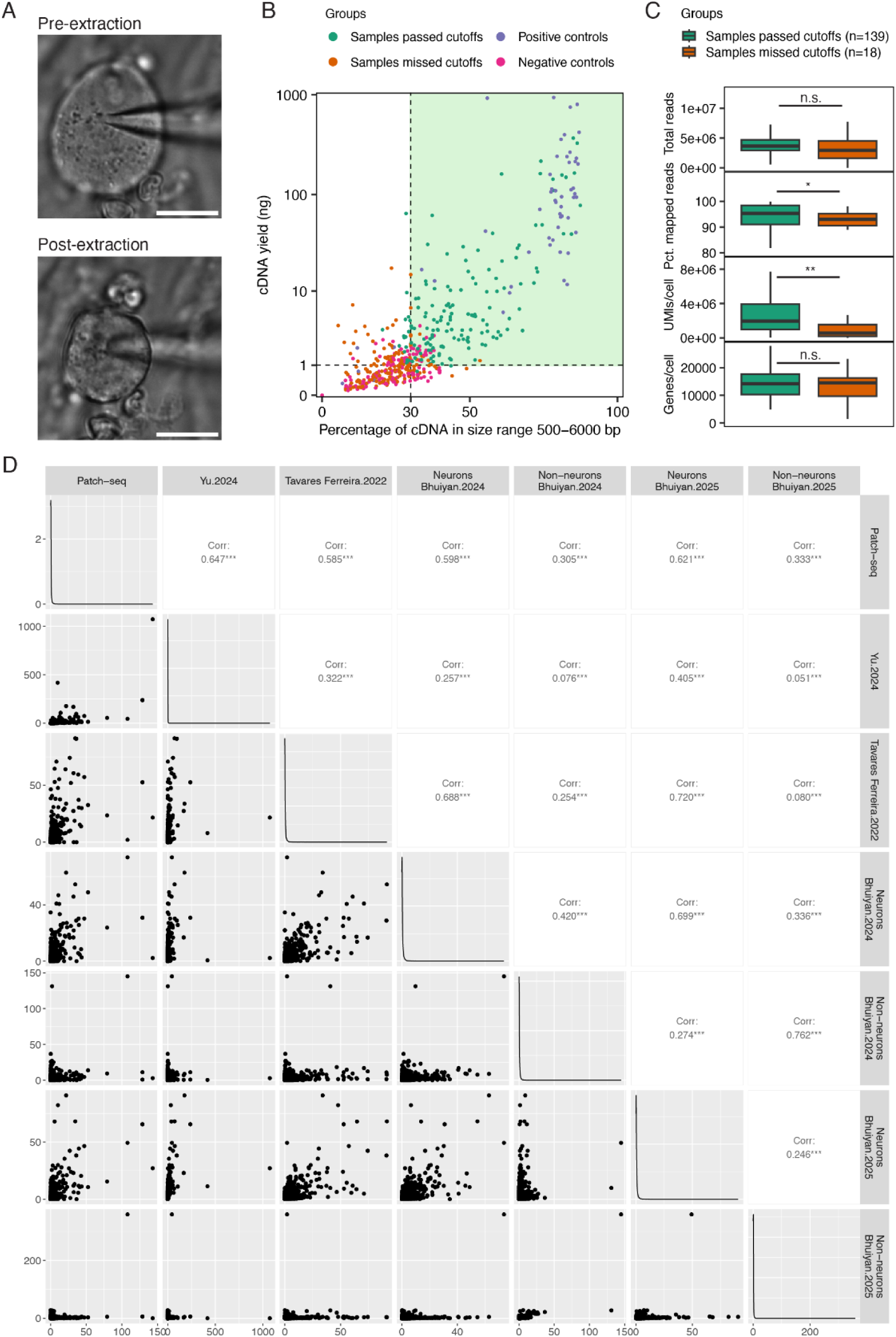
Quality control measures for hDRG Patch-seq data. (**A**) Example images of cytosolic content extraction from a recorded neuron. Shrinkage of the soma size can be confirmed visually during the extraction process. Scale bar = 30 µm. (**B**) QC metrics of the cDNA library prepared from Patch-seq samples. Samples with cDNA yield > 1 ng, at least 30% of cDNA within the size range 500-6000 bp, and both cDNA yield and cDNA percent in range are at least 25% higher than associated negative controls are considered as high quality and proceed to library preparation and sequencing. Positive controls are prepared from 50 ng of total mouse brain RNA. Negative controls are prepared from either internal solution or nuclease-free water. (**C**) Sequencing metrics of the Patch-seq samples that passed the cDNA QC (n = 139) compared to the ones that failed QC (n = 19). N.s., Not significant. Total reads: p=0.1011, two-way student T-test. Percent of mapped reads: p=0.0320, two-way student T-test. UMIs: p=0.0064, two-way student T-test. Number of genes: p=0.5522, two-way student T-test. (**D**) Pairwise correlation of hDRG Patch-seq data and published human DRG cell atlas. Bottom left panels show the normalized counts (log2CPM) of individual genes in two datasets, each averaged across all cells/nuclei in the indicated cell population from the dataset. Top right panels show the Pearson’s correlation coefficient based on the average expression of genes between dataset pairs.

**Fig S6.**
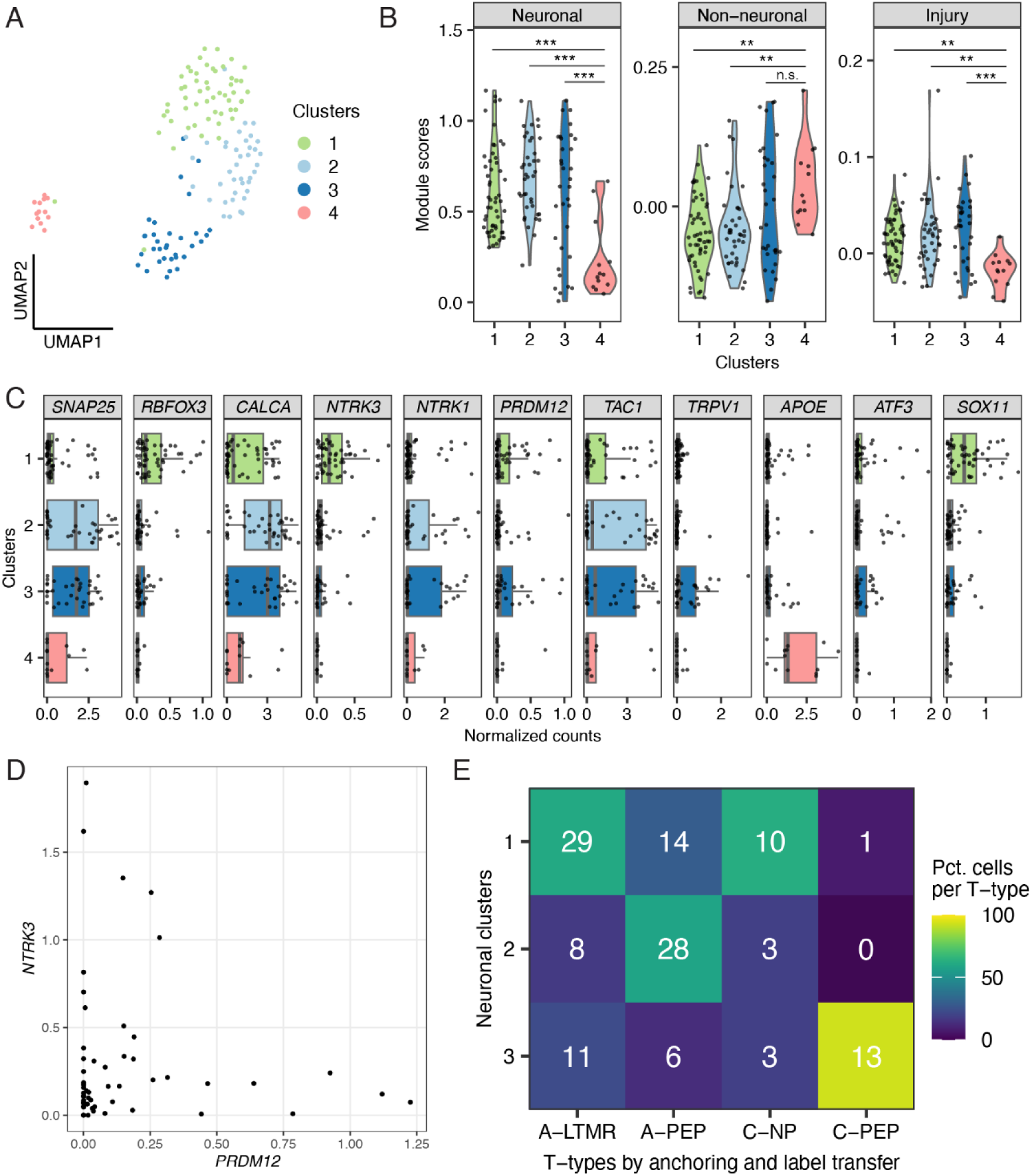
Transcriptional clustering and T-type annotation of hDRG Patch-seq samples. (**A**) UMAP of 139 Patch-seq samples, colored by transcriptional clusters. Cluster 1 (n = 54), cluster 2 (n = 39), cluster 3 (n = 33), cluster 4 (n = 13). (**B**) Violin plots showing the module scores in samples from different transcriptional clusters. Module scores were generated by aggregating the expression levels of top 200 marker genes identified from publicly available datasets (see Methods). One-way ANOVA tests, p = 3.5e-06 (***) for neuronal module score, p = 0.000491 (***) for non-neuronal module score, and p = 0.000872 (***) for injury module score. (**C**) Normalized counts (log_2_CPM) of select hDRG cell class marker genes in each transcriptional cluster. Scatter plot indicats the expression in individual samples and the boxes indicate quartiles and whiskers are 1.5-times the interquartile range (Q1-Q3). The median is a grey line inside each box. (**D**) Scatter plot showing the normalized counts (log_2_CPM) of A-LTMR transcription factor *NTRK3* and C-fiber transcription factor *PRDM12* in individual Patch-seq samples from cluster 1. (**E**) Heatmap showing the percent (as the color gradient) and cell counts (as numbers on the heatmap) of Patch-eq samples in each transcriptional neuronal cluster (clusters 1-3) that are assigned to T-types identified by anchoring to hDRG single-soma RNA-seq data.

**Fig S7.**
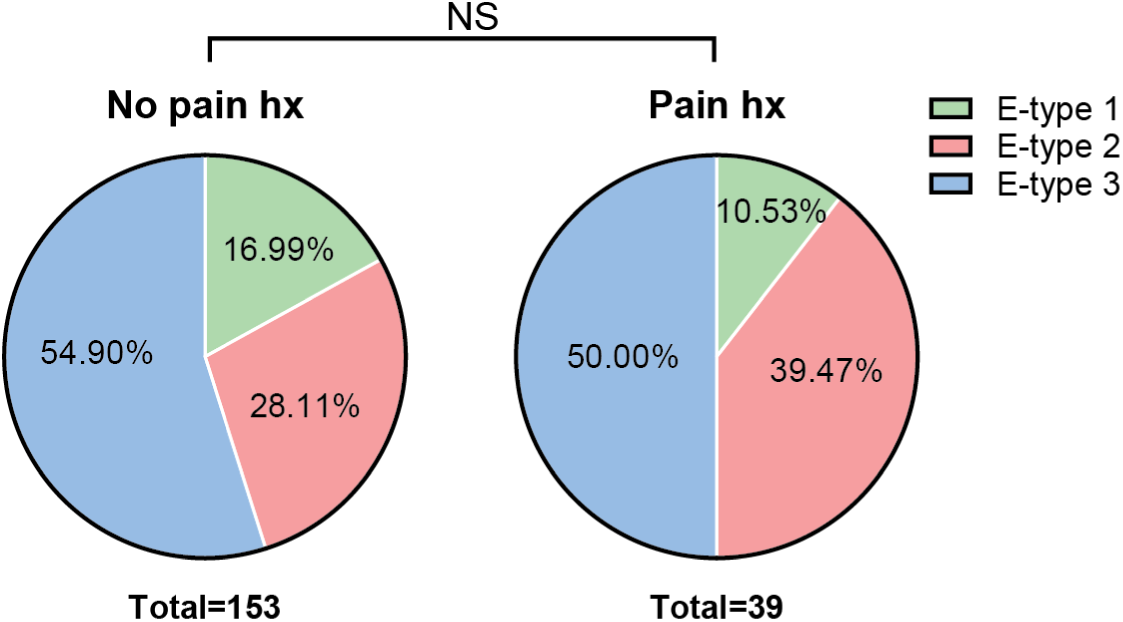
E-type distributions are not affected by pain history. Pie charts of the relative proportion of each hDRG E-type in donors with (Pain hx, right) and without (No pain hx, left) prior pain history. NS, not significant. Chi-square test, p=0.4540. N=153 neurons (No Pain Hx) and 39 neurons (Pain Hx).

**Fig S8.**
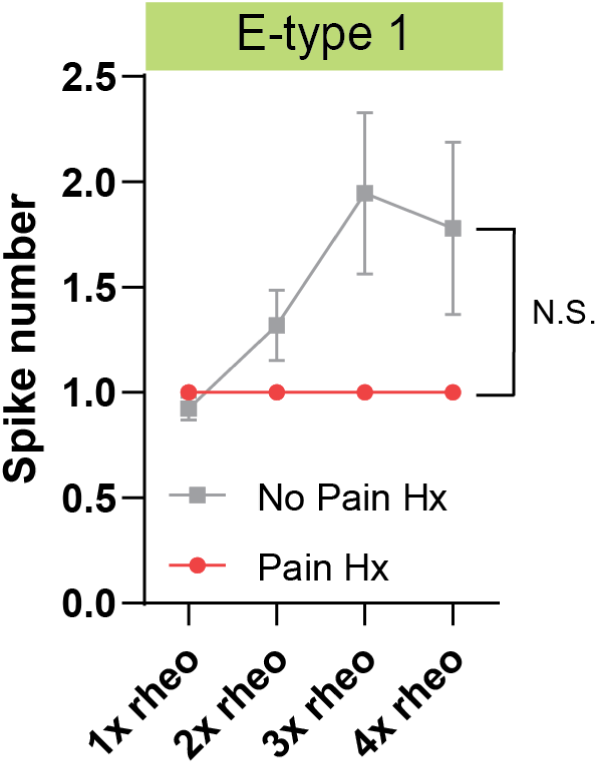
hDRG firing frequencies in E-type 1 separated by prior pain history. Input-output curves of AP frequency vs depolarizing current injections (1-4x rheobase) in hDRG with E-type 1 obtained from donors with (red; n=4 neurons) or without (grey; n=26 neurons) prior pain history. N.S., not significant. Mixed effects model analysis shows no significant fixed effect of current step (p=0.5921), pain history (p=0.2970), or interaction of current step x pain history (p=0.5921).

**Fig S9.**
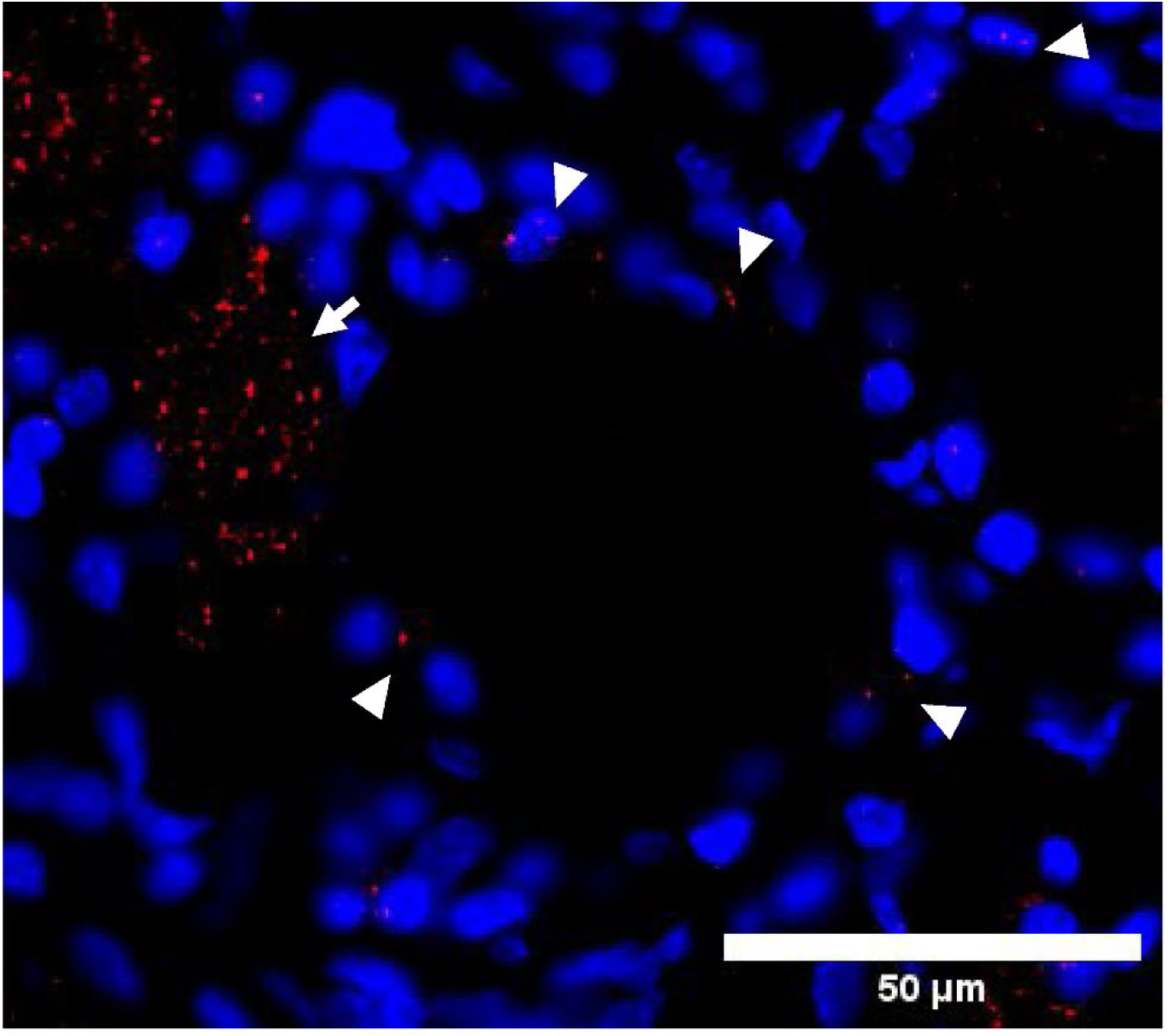
*SCN9A* expression in satellite glial cells. Arrowheads point to mRNA probes detected in satellite glial cells that surround the hDRG neuronal cell bodies. Arrow point to an adjacent neuron that expresses *SCN9A*. Blue=DAPI, red=*SCN9A* transcripts.

### Supplementary Text: PRECISION Human Pain Network group members

The members of the NIH PRECISION Human Pain Network are: Abby Pei-ting Chiu, Allison M Barry, Amy Anderson, Asta Arendt-Tranholm, Andi Wangzhou, Ayesha Ahmad, Cathryn Payne, Christoph Paul Hofstetter, Claudio Tatsui, Diana Tavares Ferreira, Felipe Espinosa, Gregory O Dussor, Ishwarya Sankaranarayanan, Jeffrey Jarvik, Joseph B Lesnak, Juan Pablo Cata, Judith A Turner, Karen Segar, Katelyn Sadler, Katherin Althea Gabriel, Khadijah Mazhar, Marisol Mancilla-Moreno, Megan L Uhelski, Michael D Burton, Michele Curatolo, Muhammad Saad Yousuf, Nguyen Tran, Olivia Catherine Davis, Patrick M Dougherty, Robert Yates North, Stephanie Shiers, Theodore J Price, Úrzula Franco-Enzástiga, William Renthal, Evangelia Semizoglou, Shamsuddin A. Bhuiyan, Parth Bhatia, Dustin Griesemer, Erika K. Williams, Jiaxiang Wang, Lily S He, Hannah MacMillan, Clifford Woolf, Barbara Gomez, Aakansksha Jain, Selwyn Jayaker, Brian Wainger, Sanghun Lee, Xianjun Dong, Weiqiang Liu, Himanshu Chintalapudi, Anthony Cicalo, Jeffrey Moffitt, Hao Zhang, James R. Stone, Iris A Lopez, Kyle Eberlin, Floris V. Raasveld, Kevin Spiegler, Wenqin Luo, Huasheng Yu, Eric A. Kaiser, Caitlin E. Cronin, Ebenezer Simpson, Hao Wu, Julie Leu, Dongming Liang, Ying Li, Mingyao Li, Hanying Yan, Patrik Ernfors, Dmitry Usoskin, Hakan Olausson, Saad Nagi, Katarina Laurell, Maria Bograkou, Otmane Bouchatta, Ewa Jarocka, Johan Nikesjo, Robert W. Gereau IV, Bryan A. Copits, Rakesh Kumar, Grace E. Moore, Deblina Nandi, Kesav Kothapalli, David M Brogan, Christopher J Dy, Lite Yang, Juliet Mwirigi, John S Del Rosario, Jun-Nan Li, Prashant Gupta, Adam Dourson, Maria Payne, Alexander Chamessian, Gary F. Marklin, John A Lemen, Elvisa Mehinovic, Zitian Tang, Jenna Ulibarri, Emma Casey, Zefan Li, Brian Yu, Sheng Chih Jin, Kevin Boyer, Ibrahim Olabayode Saliu, Bijesh George, George Murray, Huma Naz, Guoyan Zhao, Valeria Cavalli, Pauline Meriau, Sarah F. Rosen, Isabelle Gordon, Jeffrey Milbrandt, Aaron DiAntonio, Aldrin KY Yim, Amy Strickland, Liya Yuan, Joseph A Bloom, Richard Slivicki, Jyl Boline, Sam Kessler, Joost Wagenaar, Maryann Martone, Sue Tappan

