## Supplementary Figures and Tables for "Multimodal profiling of human sensory neurons links electrical properties to transcriptional identity"

**List of Supplementary Materials**

Supplementary Tables

Table S1. Detailed donor information (see separate Excel file)

Table S2. Comparison of Rheobase between hDRG neurons with different firing patterns

Table S3. DEG T-types (see separate Excel file)

Table S4. DEG E-types (see separate Excel file)

Table S5. Summary of donor demographic information, segmented by pain history

Table S6. Summary of electrophysiological features of E-type 1 between Pain Hx and No Pain Hx donors

Table S7. Summary of electrophysiological features of E-type 2 between Pain Hx and No Pain Hx donors

Table S8. Summary of electrophysiological features of E-type 3 between Pain Hx and No Pain Hx donors

Table S9. Statistics Table (see separate Excel file)

Supplementary Figures

Fig S1. Elbow plot showing the within-cluster sum of squares against different K values

Fig S2. E-type distribution across days in vitro

Fig S3. Additional membrane properties of hDRG E-types

Fig S4. Soma diameters across hDRG E-types

Fig S5. Quality control measures for hDRG Patch-seq data

Fig S6. Transcriptional clustering and T-type annotation of hDRG Patch-seq samples

Fig S7. E-type distributions are not affected by pain history

Fig S8. hDRG firing frequencies in E-type 1 separated by prior pain history

Supplementary Text: PRECISION Human Pain Network group members

**Table S2. Comparison of Rheobase between hDRG neurons with different firing patterns**

| | High Rheobase | Single | Repetitive |
|---|---|---|---|
| **N** | 30 | 97 | 101 |
| **Rheobase (Mean ± SEM)*** | 1707 ± 90.9 †,‡ | 942.2 ± 66.3 § | 511.4 ± 45.4 |

*p<0.0001, Kruskall Wallis Test
† p<0.0001, Dunn's Multiple Comparison test, High Rheobase vs. repetitive
‡ p<0.0001, Dunn's Multiple Comparison test, High Rheobase vs. Single
§ p<0.0001, Dunn's Multiple Comparison test, Single vs. Repetitive

**Table S5. Summary of hDRG donor characteristics, segmented by pain history**

| | No Pain Hx | Pain Hx |
|---|---|---|
| **N** | 34 | 13 |
| ***Race*** | | |
| **White** | 24 | 9 |
| **Black** | 9 | 3 |
| **Other/Unknown** | 1 | 1 |
| ***Sex*** | | |
| **Male** | 20 | 8 |
| **Female** | 14 | 5 |
| ***Cause of Death*** | | |
| **Anoxia** | 18 | 5 |
| **Stroke** | 11 | 5 |
| **Head Trauma** | 4 | 3 |
| **Other/Unknown** | 1 | 0 |
| **Age (years)** †,(NS) | 42.74±16.50 | 49.92±12.98 |
| **Body mass index** †, (NS) | 27.56±7.13 | 28.20±6.36 |

† Displayed in Mean ± SD
NS, Not significant; p>0.05, unpaired t-tests

**Table S6. Summary of electrophysiological features of E-type 1 between Pain Hx and No Pain Hx donors**

| | No Pain Hx (n=26 cells) | Pain Hx (n=4 cells) | P-value |
|---|---|---|---|
| **Resting membrane potential (mV)** | -49.02 ± 1.59 | -44.12 ± 5.52 | 0.46 † |
| **Input resistance (MΩ)** | 89.11 ± 8.97 | 158.1 ± 65.87 | 0.33 † |
| **Voltage sag ratio** | 0.24 ± 0.03 | 0.28 ± 0.12 | 0.69 ‡ |
| **Rheobase (pA)** | 1182.66 ± 94.98 | 1654.52 ± 176.74 | 0.0732 ‡ |
| **AP threshold (mV)** | -6.46 ± 2.18 | -10.95 ± 5.76 | 0.4597 ‡ |
| **AP Amplitude (mV)** | 27.78 ± 2.54 | 39.86 ± 4.24 | 0.0837 ‡ |
| **AP Peak (overshoot; mV)** | 21.32 ± 2.56 | 34.20 ± 11.03 | 0.10 ‡ |
| **Half-width (ms)** | 2.80 ± 0.29 | 2.73 ± 0.80 | 0.89 † |
| **Rise time (ms)** | 1.08 ± 0.10 | 1.95 ± 0.96 | 0.51 † |
| **Decay time (ms)** | 4.75 ± 0.49 | 3.16 ± 0.84 | 0.24 † |

† Mann-Whitney test
‡ Unpaired t-test

**Table S7. Summary of electrophysiological features of E-type 2 between Pain Hx and No Pain Hx donors**

| | No Pain Hx (n=43 cells) | Pain Hx (n=15 cells) | P-value |
|---|---|---|---|
| **Resting membrane potential (mV)** | -49.59 ± 2.09 | -55.78 ± 3.42 | 0.14 ‡ |
| **Input resistance (MΩ)** | 157.12 ± 18.80 | 162.63 ± 23.52 | 0.54 † |
| **Voltage sag ratio** | 0.30 ± 0.03 | 0.41 ± 0.03 | **0.0046** † |
| **Rheobase (pA)** | 323.05 ± 32.24 | 288.45 ± 46.55 | 0.87 † |
| **AP threshold (mV)** | -16.53 ± 1.25 | -19.27 ± 1.77 | 0.25 ‡ |
| **AP Amplitude (mV)** | 65.73 ± 1.58 | 74.71 ± 1.89 | **0.0022** † |
| **AP Peak (overshoot; mV)** | 49.20 ± 1.16 | 55.44 ± 1.88 | 0.0078 ‡ |
| **Half-width (ms)** | 5.73 ± 0.40 | 4.59 ± 0.47 | 0.12 † |
| **Rise time (ms)** | 0.84 ± 0.05 | 0.71 ± 0.04 | 0.16 † |
| **Decay time (ms)** | 10.11 ± 0.65 | 8.84 ± 0.80 | 0.44 † |

† Mann-Whitney test
‡ Unpaired t-test

**Table S8. Summary of electrophysiological features of E-type32 between Pain Hx and No Pain Hx donors**

| | No Pain Hx (n=84 cells) | Pain Hx (n=19 cells) | P-value |
|---|---|---|---|
| **Resting membrane potential (mV)** | -54.73 ± 0.67 | -52.44 ± 0.94 | 0.031 † |
| **Input resistance (MΩ)** | 66.29 ± 6.47 | 50.83 ± 12.85 | 0.068 † |
| **Voltage sag ratio** | 0.45 ± 0.17 | 0.53 ± 0.04 | 0.0216 † |
| **Rheobase (pA)** | 1074.41 ± 76.77 | 1175.25 ± 144.95 | 0.43 † |
| **AP threshold (mV)** | -21.86 ± 1.92 | -20.12 ± 1.92 | 0.78 † |
| **AP Amplitude (mV)** | 71.90 ± 1.22 | 71.14 ± 2.41 | 0.79 ‡ |
| **AP Peak (overshoot; mV)** | 50.46 ± 1.08 | 51.02 ± 1.74 | 0.83 ‡ |
| **Half-width (ms)** | 2.31 ± 0.12 | 1.78 ± 0.19 | 0.0458 † |
| **Rise time (ms)** | 0.73 ± 0.08 | 0.55 ± 0.05 | 0.074 † |
| **Decay time (ms)** | 3.99 ± 0.21 | 2.96 ± 0.33 | 0.034 † |

† Mann-Whitney test
‡ Unpaired t-test

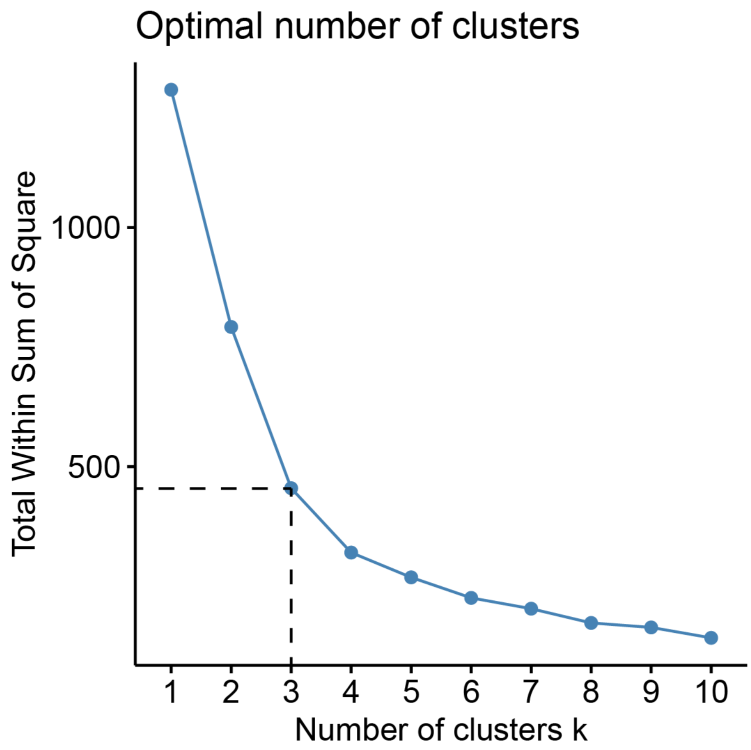


**Fig S1. Elbow plot showing the within-cluster sum of squares against different K values.** We chose k=3 as the ideal cluster number for balancing model fit and complexity.

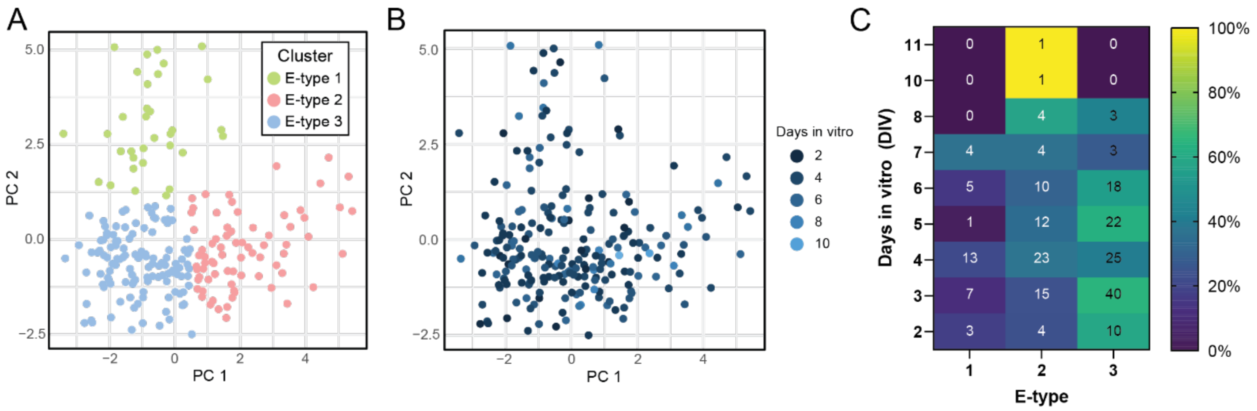


**Fig S2. E-type distribution across days in vitro.**

(**A**) Score plot of individual hDRG neurons clustered by their electrophysiological properties.

(**B**) Same plot in panel A, but each neuron is color-coded by the number of days in culture before recording.

(**C**) Heat map of E-type distributions across DIV. Number in each box shows number of cells corresponding to the E-type on the X-axis that were recorded at the DIV specified on the Y-axis. Each box is color-coded based on the relative proportion (%) of each E-type out of all the cells recorded on a particular DIV.

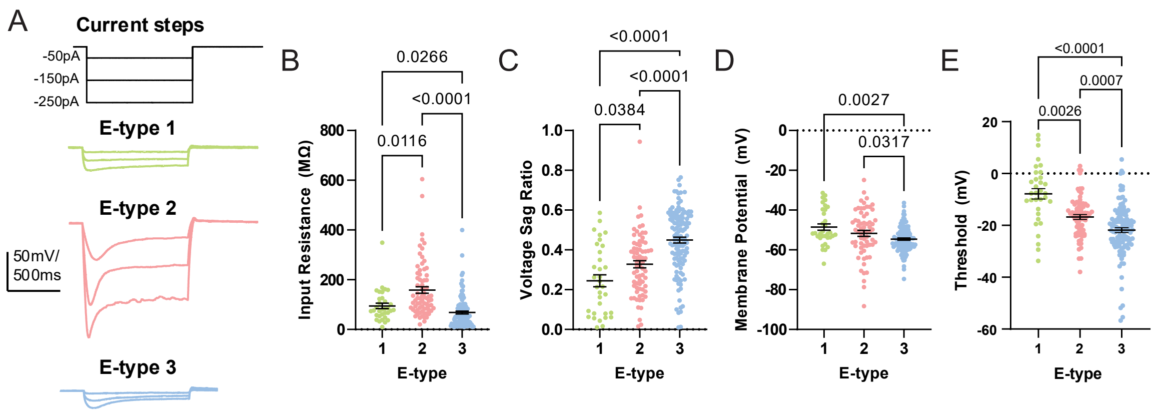


**Fig S3. Additional membrane properties of hDRG E-types.** (**A**) Example voltage traces of each hDRG E-type in response to 1 second hyperpolarizing current injection steps.
(**B-E**) Summary graphs of input resistance (**B**), voltage sag (**C**), resting membrane potential (**D**), and AP threshold (**E**) for each hDRG E-type. N=33 (E-type 1), 74 (E-type 2), and 121 neurons (E-type 3). Bonferroni correction for multiple comparisons were used to determine threshold for statistical significance (α=0.005) for group comparisons using Kruskall-Wallis or ANOVA tests. P-values for Dunn's post-hoc test (Kruskall-Wallis) or Tukey's post-hoc test (ANOVA) are shown in plot.
(**B**) Input resistance, p<0.0001 (significant), Kruskall-Wallis test.
(**C**) Voltage sag ratio, , p<0.0001 (significant), one-way ANOVA.
(**D**) Resting membrane potential, p<0.0001 (significant), Kruskall-Wallis test.
(**E**) AP threshold, p<0.0001 (significant), Kruskall-Wallis test.

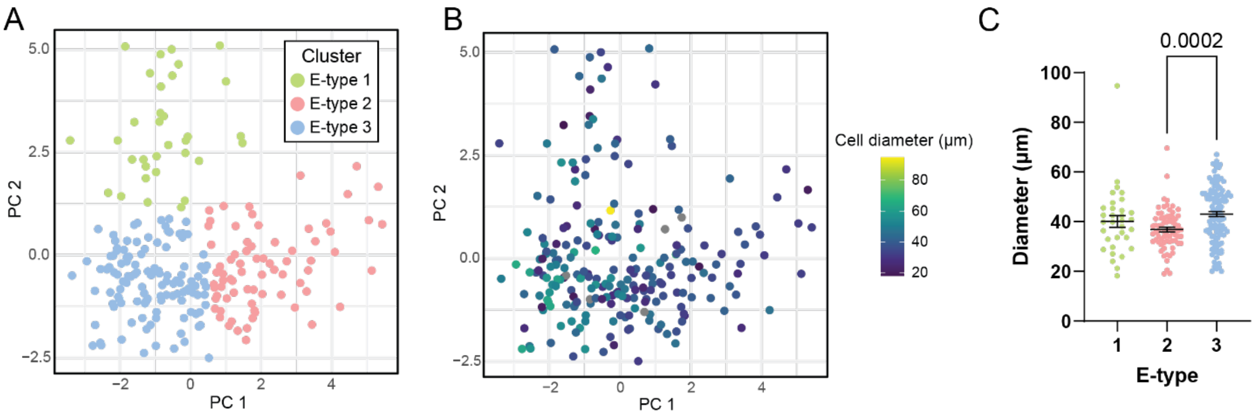


**Fig S4. Soma diameters across hDRG E-types.**

(**A**) Score plot of individual hDRG neurons clustered by their electrophysiological properties.

(**B**) Same PCA plot in (**A**) color-coded by a heat map of soma diameters in microns.

(**C**) Summary graph of soma diameter for each hDRG E-type. P=0.0003, Kruskall-Wallis test. N=33 (E-type 1), 74 (E-type 2), and 119 neurons (E-type 3). Post-hoc comparisons: p=0.0002, E-type 2 vs. E-type 3; p=0.6449, E-type 1 vs. E-type 2; p=0.2671, E-type 1 vs. E-type 3; Dunn's post-hoc test.

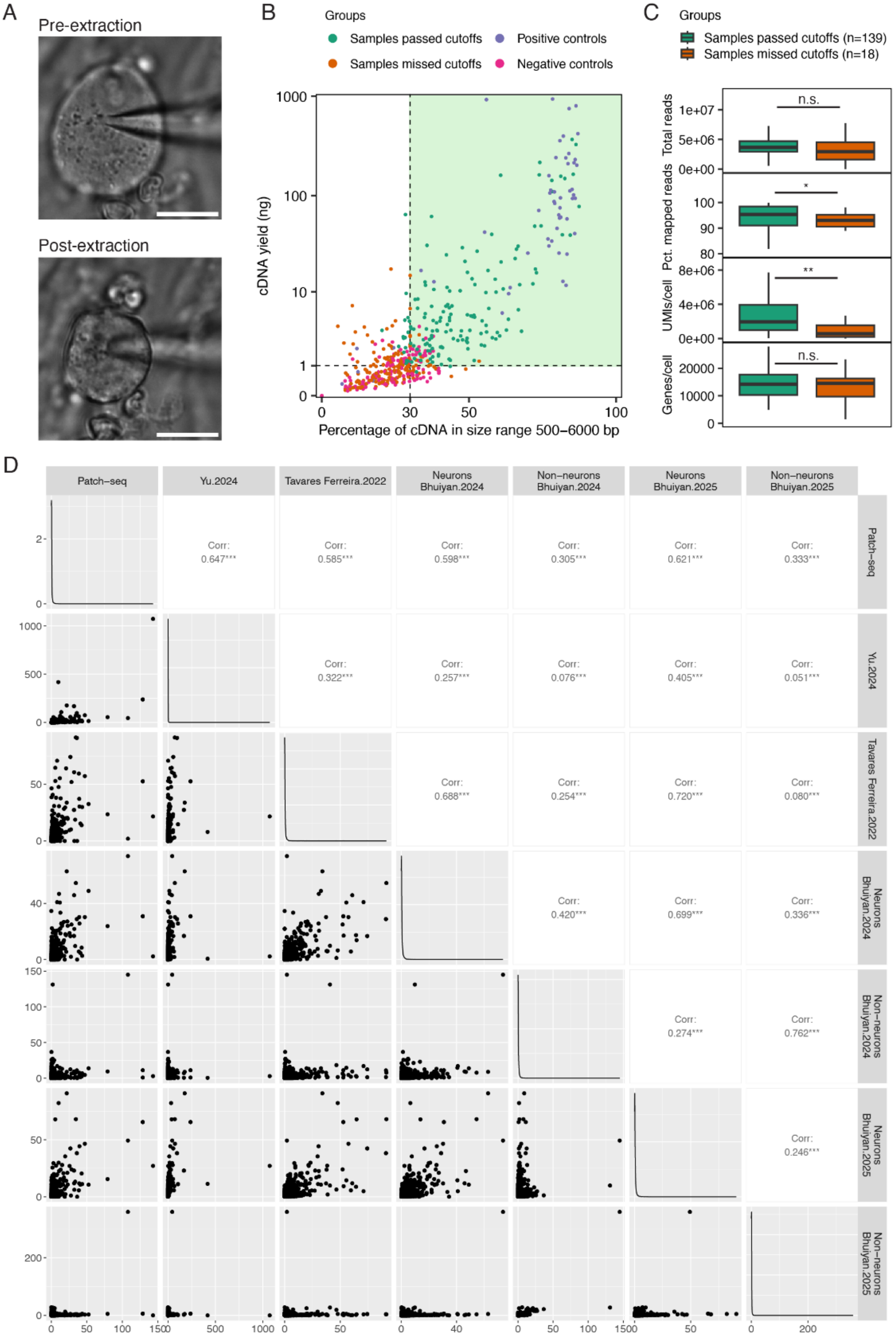

A
Pre-extraction
Post-extraction
B
Groups
Samples passed cutoffs
Positive controls
Samples missed cutoffs
Negative controls
cDNA yield (ng)
Percentage of cDNA in size range 500–6000 bp
C
Groups
Samples passed cutoffs (n=139)
Samples missed cutoffs (n=18)
Total reads
Pct. mapped reads
UMIs/cell
Genes/cell
n.s.
*
**
n.s.
D
Patch–seq
Yu.2024
Tavares Ferreira.2022
Neurons Bhuiyan.2024
Non–neurons Bhuiyan.2024
Neurons Bhuiyan.2025
Non–neurons Bhuiyan.2025
Corr: 0.647***
Corr: 0.585***
Corr: 0.598***
Corr: 0.305***
Corr: 0.621***
Corr: 0.333***
Corr: 0.322***
Corr: 0.257***
Corr: 0.076***
Corr: 0.405***
Corr: 0.051***
Corr: 0.688***
Corr: 0.254***
Corr: 0.720***
Corr: 0.080***
Corr: 0.420***
Corr: 0.699***
Corr: 0.336***
Corr: 0.274***
Corr: 0.762***
Corr: 0.246***

**Fig. S5. Quality control measures for hDRG Patch-seq data.**

(**A**) Example images of cytosolic content extraction from a recorded neuron. Shrinkage of the soma size can be confirmed visually during the extraction process. Scale bar = 30 μm.

(**B**) QC metrics of the cDNA library prepared from Patch-seq samples. Samples with cDNA yield > 1 ng, at least 30% of cDNA within the size range 500-6000 bp, and both cDNA yield and cDNA percent in range are at least 25% higher than associated negative controls are considered as high quality and proceed to library preparation and sequencing. Positive controls are prepared from 50 ng of total mouse brain RNA. Negative controls are prepared from either internal solution or nuclease-free water.

(**C**) Sequencing metrics of the Patch-seq samples that passed the cDNA QC (n = 139) compared to the ones that failed QC (n = 19). N.s., Not significant. Total reads: p=0.1011, two-way student T-test. Percent of mapped reads: p=0.0320, two-way student T-test. UMIs: p=0.0064, two-way student T-test. Number of genes: p=0.5522, two-way student T-test.

(**D**) Pairwise correlation of hDRG Patch-seq data and published human DRG cell atlas. Bottom left panels show the normalized counts (log2CPM) of individual genes in two datasets, each averaged across all cells/nuclei in the indicated cell population from the dataset. Top right panels show the Pearson's correlation coefficient based on the average expression of genes between dataset pairs.

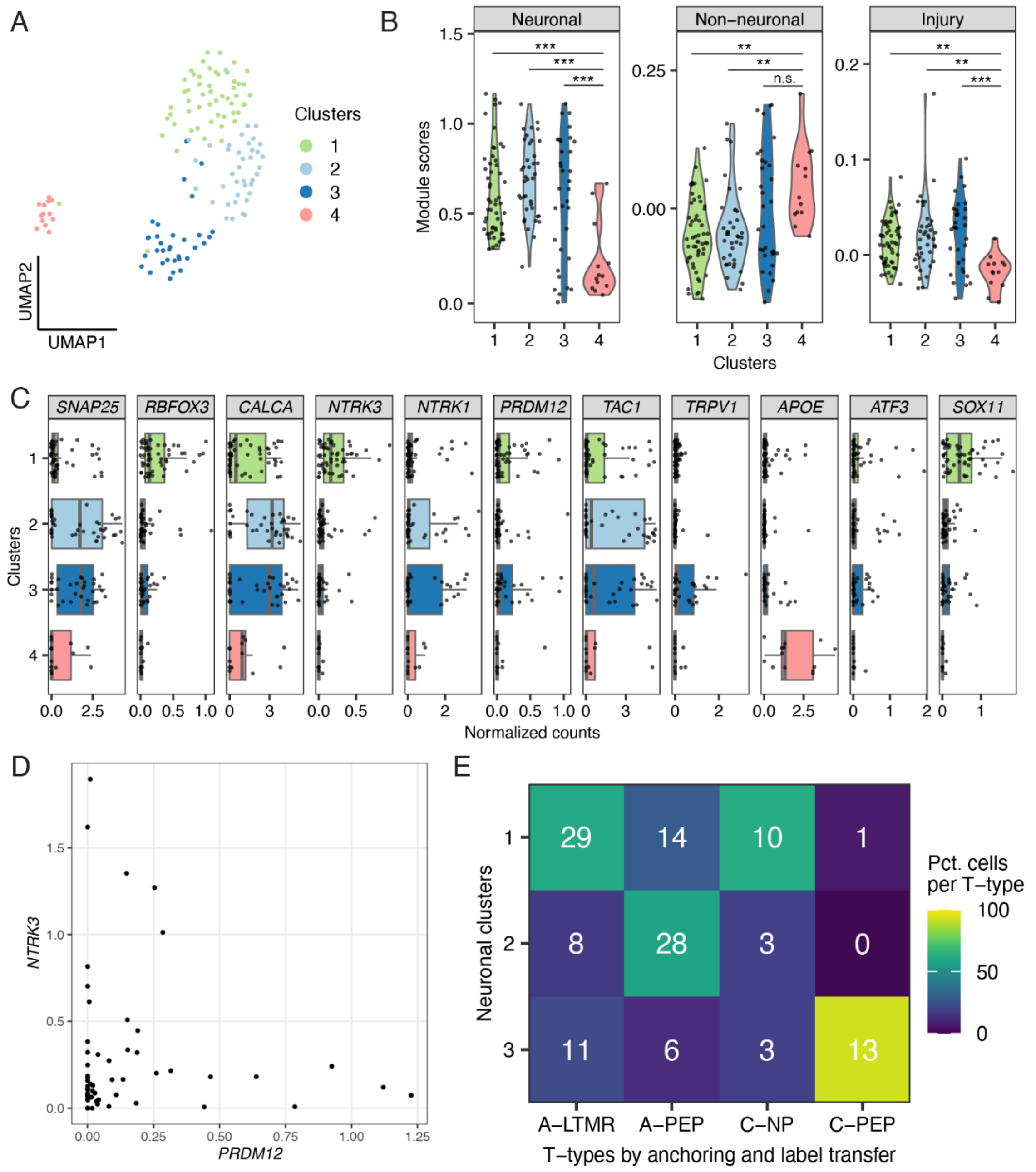


**Fig S6. Transcriptional clustering and T-type annotation of hDRG Patch-seq samples.**
(**A**) UMAP of 139 Patch-seq samples, colored by transcriptional clusters. Cluster 1 (n = 54), cluster 2 (n = 39), cluster 3 (n = 33), cluster 4 (n = 13).
(**B**) Violin plots showing the module scores in samples from different transcriptional clusters. Module scores were generated by aggregating the expression levels of top 200 marker genes identified from publicly available datasets (see Methods). One-way ANOVA tests, p = 3.5e-06 (***) for neuronal module score, p = 0.000491 (***) for non-neuronal module score, and p = 0.000872 (***) for injury module score.

(**C**) Normalized counts ($log_2CPM$) of select hDRG cell class marker genes in each transcriptional cluster. Scatter plot indicats the expression in individual samples and the boxes indicate quartiles and whiskers are 1.5-times the interquartile range (Q1-Q3). The median is a grey line inside each box.
(**D**) Scatter plot showing the normalized counts ($log_2CPM$) of A-LTMR transcription factor *NTRK3* and C-fiber transcription factor *PRDM12* in individual Patch-seq samples from cluster 1.
(**E**) Heatmap showing the percent (as the color gradient) and cell counts (as numbers on the heatmap) of Patch-eq samples in each transcriptional neuronal cluster (clusters 1-3) that are assigned to T-types identified by anchoring to hDRG single-soma RNA-seq data.

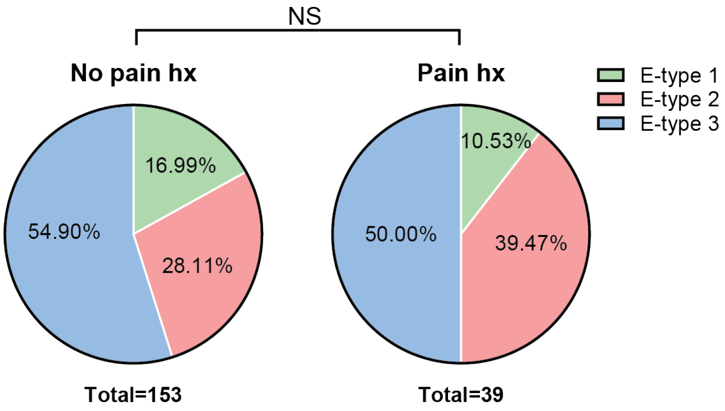


**Fig S7. E-type distributions are not affected by pain history.** Pie charts of the relative proportion of each hDRG E-type in donors with (Pain hx, right) and without (No pain hx, left) prior pain history. NS, not significant. Chi-square test, p=0.4540. N=153 neurons (No Pain Hx) and 39 neurons (Pain Hx).

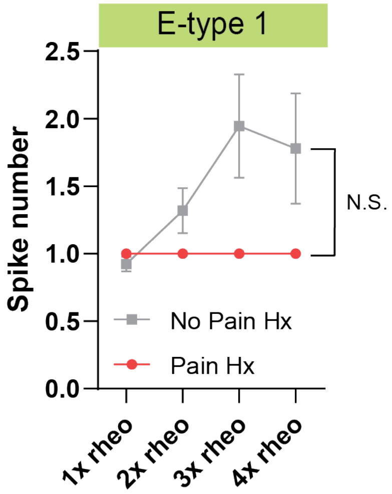


**Fig S8. hDRG firing frequencies in E-type 1 separated by prior pain history.** Input-output curves of AP frequency vs depolarizing current injections (1-4x rheobase) in hDRG with E-type 1 obtained from donors with (red; n=4 neurons) or without (grey; n=26 neurons) prior pain history. N.S., not significant. Mixed effects model analysis shows no significant fixed effect of current step (p=0.5921), pain history (p=0.2970), or interaction of current step x pain history (p=0.5921).

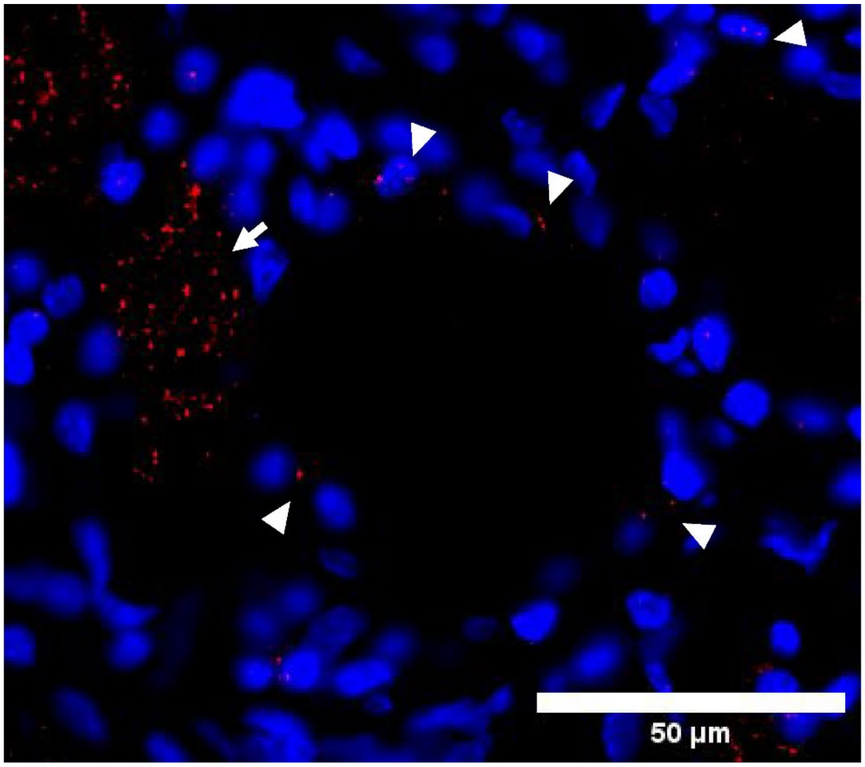


**Fig S9.** ***SCN9A*** **expression in satellite glial cells.** Arrowheads point to mRNA probes detected in satellite glial cells that surround the hDRG neuronal cell bodies. Arrow point to an adjacent neuron that expresses *SCN9A*. Blue=DAPI, red=*SCN9A* transcripts.
